# Modelling the contributions to hyperexcitability in a mouse model of Alzheimer’s disease

**DOI:** 10.1101/2022.06.29.494500

**Authors:** Martin Mittag, Laura Mediavilla, Stefan Remy, Hermann Cuntz, Peter Jedlicka

## Abstract

Neuronal hyperexcitability is a feature of Alzheimer’s disease (AD). Three main mechanisms have been proposed to explain it: i), dendritic degeneration leading to increased input resistance, ii), ion channel changes leading to enhanced intrinsic excitability, and iii), synaptic changes leading to excitation-inhibition (*E/I*) imbalance. However, the relative contribution of these mechanisms is not fully understood. Therefore, we performed biophysically realistic multi-compartmental modelling of excitability in reconstructed CA1 pyramidal neurons of wild-type and APP/PS1 mice, a well-established animal model of AD. We show that, for synaptic activation, the excitability promoting effects of dendritic degeneration are cancelled out by excitability decreasing effects of synaptic loss. We find an interesting balance of excitability regulation with enhanced degeneration in the basal dendrites of APP/PS1 cells potentially leading to increased excitation by the apical but decreased excitation by the basal Schaffer collateral pathway. Furthermore, our simulations reveal that three additional pathomechanistic scenarios can account for the experimentally observed increase in firing and bursting of CA1 pyramidal neurons in APP/PS1 mice. Scenario 1: increased excitatory burst input; scenario 2: enhanced *E/I* ratio and scenario 3: alteration of intrinsic ion channels (*I_AHP_* down-regulated; *I_Nap_*, *I_Na_* and *I_CaT_* up-regulated) in addition to enhanced *E/I* ratio. Our work supports the hypothesis that pathological network and ion channel changes are major contributors to neuronal hyperexcitability in AD. Overall, our results are in line with the concept of multi-causality and degeneracy according to which multiple different disruptions are separately sufficient but no single disruption is necessary for neuronal hyperexcitability.

**In brief:** Using a computational model, we find that changes in the extrinsic network and intrinsic biophysical neuronal properties rather than dendritic degeneration alone explain the altered firing behaviour observed in Alzheimer’s disease (AD).

**Highlights:**

- Simulations of synaptically driven responses in PCs with AD-related dendritic degeneration.
- Dendritic degeneration alone alters PC responses to layer-specific input but additional pathomechanistic scenarios are required to explain neuronal hyperexcitability in AD.
- Possible scenario 1: Burst hyperactivity of the surrounding network can explain hyper-excitability of PCs during AD.
- Possible scenario 2: AD-related increased excitatory input together with decreased inhibitory input (*E/I* imbalance) can lead to hyperexcitability in PCs.
- Possible scenario 3: Changes in *E/I* balance combined with altered ion channel properties can account for hyperexcitability in AD.

## Introduction

Neuronal hyperexcitability has been described as a characteristic feature of Alzheimer’s disease (AD, Palop and Mucke, 2009; Vossel *et al*., 2017; Zott *et al*., 2018; Kazim *et al*., 2021). It is observed in the early phases of the disease progression (Dickerson *et al*., 2005; Busche and Konnerth, 2015) at the circuit as well as single cell level. Observations of hyperexcitability in AD patients (Palop *et al*., 2007; Vossel *et al*., 2013; Palop and Mucke, 2016; Vossel *et al*., 2016; Horvath *et al*., 2021; Ranasinghe *et al*., 2022; Vossel *et al*., 2021) are consistent with data from mouse models of AD (Busche *et al*., 2008, 2012, 2015a,b; Rudinskiy *et al*., 2012; Grienberger *et al*., 2012; Scala *et al*., 2015; Maier *et al*., 2014; Šišková *et al*., 2014; Hall *et al*., 2015; Xu *et al*., 2015; Liebscher *et al*., 2016; Keskin *et al*., 2017; Müller *et al*., 2021). Most studies showing AD-associated hyperactivity such as increased frequency of calcium transients or the occurence of hyperactive neuron populations in mice (Busche *et al*., 2008, 2012; Busche and Konnerth, 2015) and rats (Sosulina *et al*., 2021) and increased seizure activity during electroencephalographic (EEG) recordings in the mouse hippocampus (Palop *et al*., 2007; Palop and Mucke, 2009) do not or can not determine if the mode of increased neuronal excitability comes from an enhanced single spike rate or from a switch to enhanced burst firing. However, the change of firing mode towards stronger burst firing is a feature in several neurological disorders such as epilepsy (Sanabria *et al*., 2001; Wellmer *et al*., 2002; Pothmann *et al*., 2019) and chronic stress (Okuhara and Beck, 1998). In AD, amyloid-beta accumulation has been linked to a change in burst firing pattern (Chen, 2005; Minkeviciene *et al*., 2009; Kellner *et al*., 2014). For example in APP/PS1 model mice, *in vivo* and *in vitro* patch-clamp and *in vivo* extracellular recordings revealed hyperactivity of CA1 pyramidal neurons in the form of increased mean firing rate as well as enhanced bursting (Šišková *et al*., 2014).

Three prominent explanations of neuronal hyperexcitability in AD (including the enhanced bursting) have been proposed (Ferrao Santos *et al*., 2010; Zott *et al*., 2018; Vyas *et al*., 2020; Maestú *et al*., 2021), which will be introduced in detail below: alterations of intrinsic properties by (i) dendritic degeneration, or by (ii) ion channel changes, or alterations of extrinsic network properties by (iii) enhanced synaptic excitation-inhibition (*E/I*) ratio.

i. Atrophic degeneration of neuronal dendrites is one of the hallmarks of AD (Braak and Braak, 1991; Braak *et al*., 1993; Anderton *et al*., 1998), well documented both in patients (Augustinack *et al*., 2002; Grutzendler *et al*., 2007; Merino-Serrais *et al*., 2013) as well as in animal models (Grutzendler *et al*., 2007; Le *et al*., 2001; Tsai *et al*., 2004; Moolman *et al*., 2004). It progressively affects brain areas that play important roles in learning and memory, such as the dentate gyrus, the CA1 and the subiculum area of the hippocampus and cerebral cortex (Spires and Hyman, 2004; Grutzendler *et al*., 2007; Adlard and Vickers, 2002; Falke *et al*., 2003; Geula *et al*., 1998). Dendritic changes have been suggested to play a major role in the pathogenesis of AD (Cochran *et al*., 2014). However, the functional consequences of dendritic degeneration associated with a concurrent synapse loss (Masliah *et al*., 1994), which is another hallmark of AD (Terry *et al*., 1991), have only recently started being elucidated in APP/PS1 mice (Šišková *et al*., 2014). The dendritic degeneration in CA1 PCs has been proposed to contribute to their hyperexcitability in the form of higher firing rates associated with enhanced bursting (Šišková *et al*., 2014). However, if also synaptic loss is considered then dendritic degeneration with its decrease in input conductance might counteract the lower number of synapses, homeostatically maintaining normal excitability (Platschek *et al*., 2016, 2017; Cuntz *et al*., 2021). To explore this possibility, we implemented both synapse loss and dendritic degeneration (Šišková *et al*., 2014) in compartmental CA1 PC models and analysed their synaptically driven activity.
ii. Modifications of intrinsic excitability due to changes in ionic channels have also been implicated in the pathogenesis of the AD (Kerrigan *et al*., 2014). Experimental studies have reported alterations in the density of several active membrane channels, such as A-type *K*^+^ channel, voltage-dependent *Na*^+^ channel, and delayed-rectifier *K*^+^ channel (Good *et al*., 1996; Kim *et al*., 2007; Brown *et al*., 2011; Scala *et al*., 2015; Liu *et al*., 2015; Wang *et al*., 2016; Ghatak *et al*., 2019). Several studies have provided evidence supporting the contribution of the reduced A-type *K*^+^ current to the hyperexcitability observed in AD-affected neurons (Chen, 2005; Morse *et al*., 2010; Culmone and Migliore, 2012; Scala *et al*., 2015; Frazzini *et al*., 2016; Rodrigues *et al*., 2017). Likewise, increased excitability has also been documented in a mouse model of AD, where it was attributed to changes in the dendritic tree and alterations in the expression and function of A-type *K*^+^ channels (Hall *et al*., 2015). There has been contradictory evidence showing the role of the hyperpolarisation-activated *H* channel with studies indicating both a decrease or an increase of its density (Musial *et al*., 2018; Vitale *et al*., 2021). Further experimental findings have revealed the influence of the small and large calcium-activated *K*^+^ channels in AD model mice (Beck and Yaari, 2008; Zhang *et al*., 2014; Wang *et al*., 2015a,b). Also the disruption of *Ca*^2+^ signaling and *Ca*^2+^ channels plays an important role in the pathogenesis of AD (Bezprozvanny and Mattson, 2008; Bojarski *et al*., 2008; Anekonda *et al*., 2011; Tan *et al*., 2012). For AD-related pathologies the L-type *Ca*^2+^ channel (Anekonda *et al*., 2011; Berridge, 2014), the A-type *K*^+^ channel (Chen, 2005) and *Na*^+^ channels (Wang *et al*., 2016; Ghatak *et al*., 2019; Müller *et al*., 2021) have been shown to be involved in burst rate amplification. Modelling studies (Medlock *et al*., 2018; Garg *et al*., 2021) confirm the role of *Ca*^2+^ channels for enhanced burst firing. Evidently, the modification of intrinsic excitability due to alterations in ion channel expression is well documented in AD. However, its interplay with synaptic and dendritic changes has not yet been fully clarified.
iii. Several studies have provided evidence for enhanced glutamatergic excitation (Busche and Konnerth, 2016; Zott *et al*., 2019) and impaired inhibition during AD (Busche *et al*., 2008, 2012; Takahashi *et al*., 2010; Schmid *et al*., 2016; Palop and Mucke, 2016; Ambrad Giovannetti and Fuhrmann, 2019; Xu *et al*., 2020; Gervais *et al*., 2021). This includes decreased perisomatic inhibition (Verret *et al*., 2012) which can alter the firing pattern towards more bursts (Pouille and Scanziani, 2004). The excitatory drive constitutes one of the main determinants of neuronal spontaneous firing rate (Frere and Slutsky, 2018). Therefore, a shift towards synaptic excitation (Roberson *et al*., 2011) that increases the *E/I* ratio may explain enhanced firing rates in AD during spontaneous activity. Epileptiform disruption of spontaneous neuronal activity in hippocampal circuits is a typical feature in mouse models of AD (Palop *et al*., 2007), where sharp synchronous discharges linked to memory deficits have been observed (Born *et al*., 2014). In line with the relevance of enhanced *E/I* ratio, recent clinical observations suggest that pharmacological suppression of glutamatergic excitation (by levetiracetam) is a promising way of improving cognition in AD patients with epileptiform hyperexcitability (Vossel *et al*., 2021).

Although these three groups of mechanisms have been proposed to account for AD-related hyperexcitability, their contributions and mutual interplay is not fully understood. Therefore, in the present study, we took advantage of the unique feature of biophysical modelling that enables the investigation of isolated and combined parameter changes. The computational approach allowed us to disentangle which changes and their contributions to hyperexcitability in AD are most relevant. First, we investigated whether the AD-related dendritic degeneration in APP/PS1 mice by itself supports hyperactivation of their CA1 pyramidal neurons or, alternatively, whether it compensates for the loss of synapses and helps maintain unchanged neuronal spike rates. Our biophysically detailed modelling of realistic dendrite morphologies from APP/PS1 mice indicates that the observed morphological changes alone do not lead to the hyperexcitability of CA1 pyramidal neurons for whole cell distributed synaptic inputs. Increased degeneration in the basal dendrites of APP/PS1 cells can lead to a region-dependent change in excitation as seen in clustered stimulations of main layer-specific CA1 input pathways. Second, in line with a multi-causal pathogenesis, we showed that the increased excitability of CA1 pyramidal cells can be sufficiently accounted for by a shift in *E/I* balance towards glutamatergic network excitation (increased network burst activity and decreased inhibitory inputs) or by its combination with changes in ion channel expression (*I_AHP_* channel density down-regulated; *I_Nap_*, *I_Na_* and *I_CaT_* up-regulated). Our results are in line with the concept of degeneracy (Tononi *et al*., 1999; Edelman and Gally, 2001), according to which similar physiological but also similar pathological states such as neural hyperexcitability can emerge from multiple structurally distinct mechanisms (Neymotin *et al*., 2016; Ratté and Prescott, 2016; O’Leary, 2018; Kamaleddin, 2022; Medlock *et al*., 2022; Stö ber *et al*., 2022).

## Results

### Dendritic degeneration alone does not account for synaptically driven hyperexcitability in APP/PS1 CA1 pyramidal cell models

Dendritic degeneration is a prominent feature observed in AD (Baloyannis, 2009). However, its functional consequences have not yet been fully understood. Investigating whether dendritic degeneration alone is sufficient to account for the pathologically increased neuronal excitability observed in AD (Vossel *et al*., 2017) is only possible with computational models that allow for isolated manipulations of morphological parameters. Therefore, we implemented data-driven, biophysically and anatomically realistic compartmental models of wild-type (WT) and APP/PS1 CA1 pyramidal neurons. We examined the impact of dendritic morphology on the cell’s output behaviour by using a previously published morphological data set of 31 WT and 28 APP/PS1 CA1 pyramidal neurons from a mouse model of AD (Šišková *et al*., 2014). Morphological data were used to create neuronal models in the simulation environment T2N (Beining *et al*., 2017). A representative morphology from each cell group is shown in **Figure 1A**.

**Figure 1.**
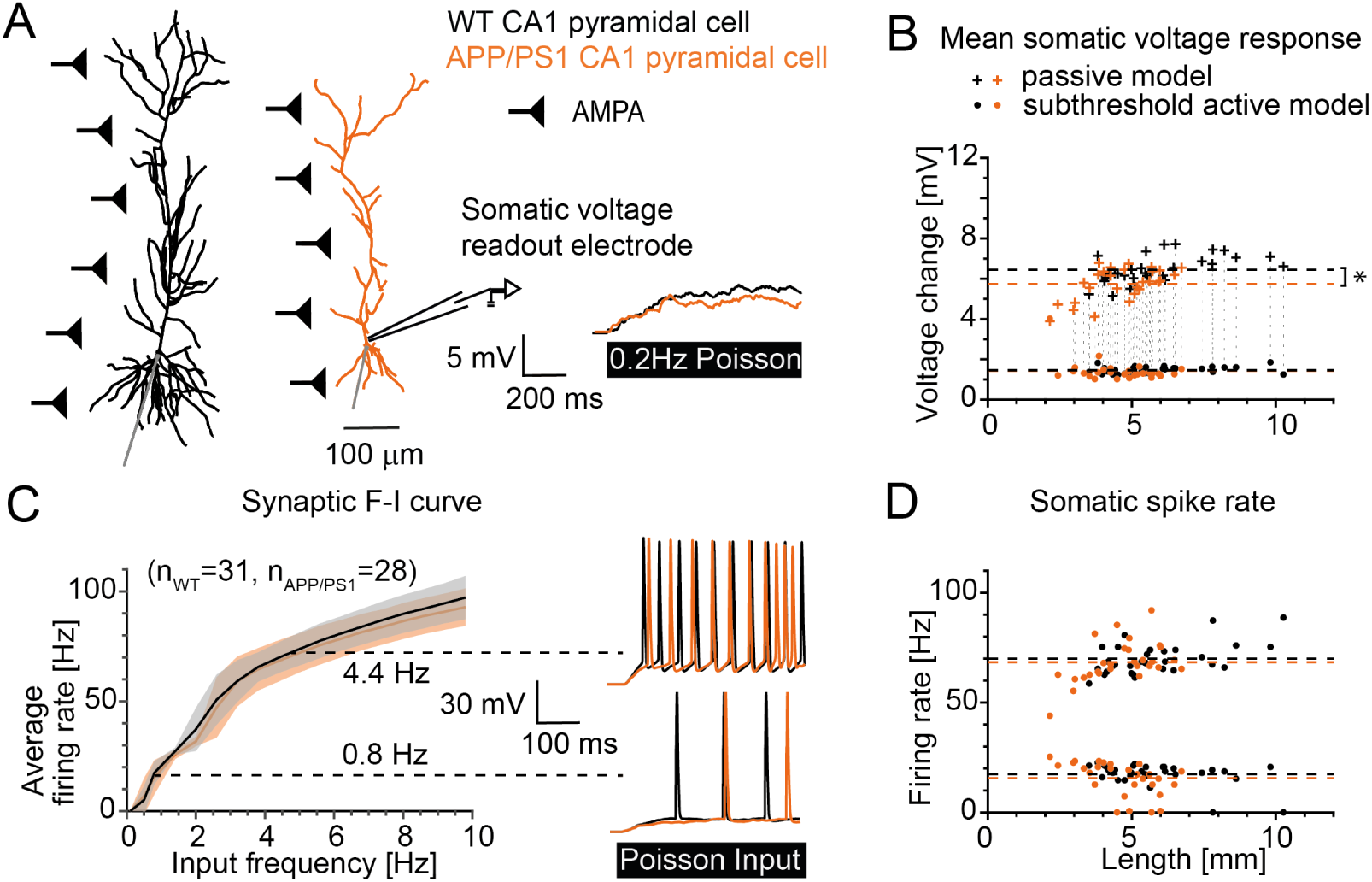
Responses to distributed synaptic AMPA inputs show that dendritic degeneration by itself causes no hyperexcitability in APP/PS1 model cells if synaptic density remains preserved. **A**, Sample 3D-reconstructed morphologies of wildtype (WT, *black*) and APP/PS1 (*orange*) CA1 pyrami-dal cells. Distributed AMPA synapses (*black* triangles) receiving Poisson input patterns. The inset on the *right* shows sample trajectories for the voltage response of the two sample cells for the passive model with synaptic stimulation at 0.2*Hz*. **B**, Voltage responses to distributed AMPA inputs at 0.2*Hz* for the passive (*crosses*) and the subthreshold active model by Poirazi *et al*. (2003b, *circles*) for all available cell morphologies (WT: *n* = 31, APP/PS1: *n* = 28). The dashed lines show the mean activity of the WT (*black*) and APP/PS1 (*orange*) CA1 cell groups (mean voltage passive model: WT 6.45 *±* 0.67*mV*, APP/PS1 5.74 *±* 0.80*mV*, the asterisk depicts *p <* 0.0005; mean voltage subthreshold active model: WT 1.47 *±* 0.17*mV*, APP/PS1 1.41 *±* 0.51*mV*). **C**, Synaptic input-output (IO) curve for the average somatic firing rate of all WT and APP/PS1 cells in active compartmental models with suprathreshold frequencies of AMPA synapse activation. The input frequency ranged from 0.1*Hz* to 10*Hz*. The *right* insets show action potential (AP) firing traces of the two sample cells for an input frequency of 0.8*Hz* and 4.4*Hz*, respectively. **D**, Firing rate versus dendritic length corresponding to the data points with input frequency of 0.8*Hz* and 4.4*Hz* in C. The dashed lines show the mean firing rate (0.8*Hz* input: WT 17.35 *±* 5.29*Hz*, APP/PS1 15.40 *±* 7.72*Hz*; 4.4*Hz* input: WT 69.91 *±* 7.23*Hz*, APP/PS1 68.29 *±* 9.41*Hz*). For all simulations the AMPA synapse strength was 0.1*nS* with rise time constants of *τ_rise_* = 0.2*ms* and decay time constants of *τ_decay_* = 2.5*ms* and the density was 1 synapse per 2*µm*.

To simulate *in vivo*-like conditions, where a single cell integrates the synaptic inputs from different locations across the whole dendritic tree, we uniformly distributed excitatory (AMPA) synapses at a fixed density along the cell’s dendritic arbour and induced Poisson-like synaptic background noise, while measuring the cell’s response at the soma (**Figure 1A**). Importantly, in agreement with the experimental spine density data from the reconstructed morphologies (Šišková *et al*., 2014), we used the same density of synapses for WT and APP/PS1 morphologies. Identical density of synapses in WT and APP/PS1 dendrites results in a smaller absolute number of synapses in shorter (degenerated) APP/PS1 dendrites. In this simple implicit way we implemented the synapse loss in APP/PS1 mice (but see also below more detailed simulations of synapse density and **Supplementary Table S3**).

Firstly, we performed simulations in electrically passive dendrites. We found that the dendritic degeneration observed in the APP/PS1 morphology group (see **Figure S2A**), did not lead to an increase in the somatic voltage responses (**Figure 1B**, cross markers), even though the reduction in cell size increases the cell’s input resistance. In fact, the APP/PS1 cell group showed on average a reduced voltage change when compared to the WT group (mean voltage passive model: WT 6.45 *±* 0.67*mV*, APP/PS1 5.74 *±* 0.80*mV*, the asterisk depicts *p <* 0.0005).

Secondly, in order to explore neuronal excitability in more realistic active dendrites, we extended the passive models by the insertion of active ion channels from a well established biophysical CA1 pyramidal cell model (Poirazi *et al*. (2003b), see Table S1 and **Supplementary Figure S1** for model details and for comparison with a second CA1 model Jarsky *et al*. (2005) in **Supplementary Figure S3**). To achieve this, we used a previously established scaling method for transferring distance-dependent ion channel densities to multiple CA1 pyramidal cell morphologies (Cuntz *et al*., 2021). The addition of the active ion channels led to a pronounced change in the subthreshold behaviour of neurons in response to synaptic activation, reducing and equalising the voltage responses across both morphology groups (**Figure 1B**, circle markers, mean voltage subthreshold active model: WT 1.47 *±* 0.17*mV*, APP/PS1 1.41 *±* 0.51*mV*). Similarly, the spiking behaviour was comparable across both cell groups, displaying overlapping *FI*-curves (**Figure 1C**) and firing rates that were independent from the cell’s dendritic length and complexity (**Figure 1D**, 0.8*Hz* input: WT 17.35 *±* 5.29*Hz*, APP/PS1 15.40 *±* 7.72*Hz*; 4.4*Hz* input: WT 69.91 *±* 7.23*Hz*, APP/PS1 68.29 *±* 9.41*Hz*). This remained true for other morphological parameters such as branch points and surface area (**Figure S2**).

Together these results show that the degenerated dendrites in the APP/PS1 cells do not lead to a facilitation of synaptic integration for either the passive or active model, when compared to their WT counterparts (see **Figure S3** for similar results in a different biophysical CA1 pyramidal cell model by Jarsky *et al*., 2005). We conclude that morphological changes alone cannot account for the cellular hyperexcitability observed in the APP/PS1 mouse model.

### Dendritic degeneration can affect CA1 PC excitability by selective gating of layer-specific inputs

Degeneration can disproportionally affect some dendritic subregions. For example, in CA1 APP/PS1 morphologies, basal dendrites were more affected than apical dendrites (Šišková *et al*., 2014). Therefore, we explored whether subregion-specific dendritic degeneration leads to a change in CA1 PC behaviour in response to layer-specific input pathways. For this purpose, we have implemented a more realistic distribution of synaptic inputs based on anatomical synaptic data. We incorporated layer-specific synapse densities as well as distance-dependent and lognormal synaptic weight distributions (based on Magee and Cook (2000); Megías *et al*. (2001); Katz *et al*. (2009); Šišková *et al*. (2014); Kim *et al*. (2015); Bloss *et al*. (2016)), which included AMPA, NMDA and GABA synapses (see **Methods** for details, **Supplementary Table S3** and **Figure 2A**). We divided the inputs into the three major layer-specific input pathways to the CA1 cell (**Figure 2C**, *Schematics*): the perforant path and the apical and basal Schaffer collaterals. Synaptic background activity (Poisson-like at 0.5*Hz*), targeting all dendritic regions, did not lead to increased excitability in the APP/PS1 cells. Moreover, the spiking behaviour still remained independent from the cell’s dendritic length (**Figure 2B**, WT 25.24 *±* 6.74*Hz*, APP/PS1 23.69 *±* 3.87*Hz*; see comparison with **Figure 1D**).

**Figure 2.**
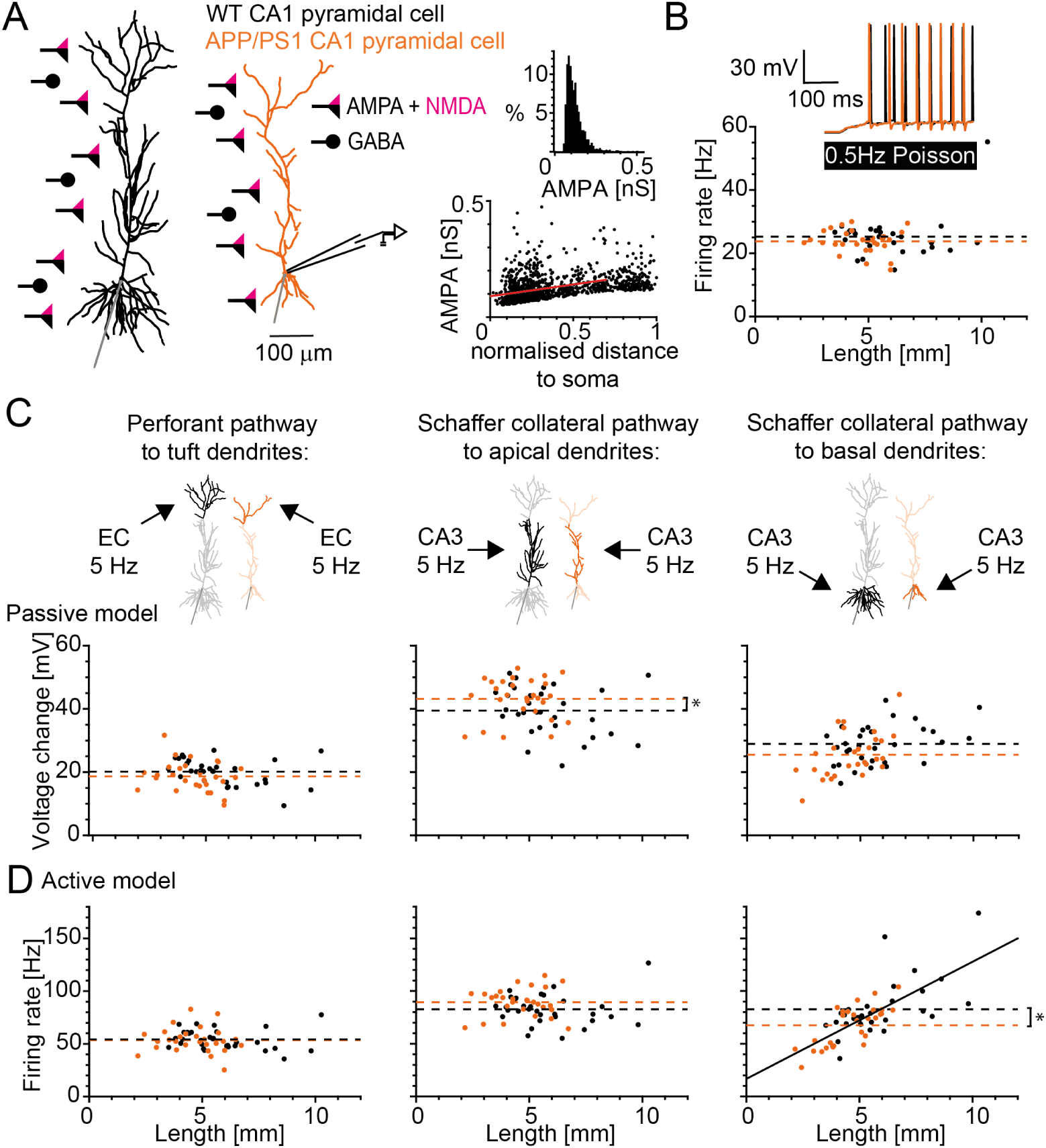
Responses to activated AMPA, NMDA and GABA synapses of main layer-specific input pathways show that dendritic degeneration by itself can cause changes in excitability in APP/PS1 model cells through selective gating of the inputs. **A**, Sample morphologies of CA1 pyramidal cells (WT in *black*, APP/PS1 in *orange*) with a schematic of distributed AMPA (*black triangle*), NMDA (*magenta triangle*) and GABA (*black circle*) synapses (**Supplementary Table S3**). *Right*, Biologically realistic distribution of AMPA weights with respect to the relative distance to soma (lognormal distribution of synaptic weights with a distance-dependent increase). **B**, Firing rate versus dendritic length of WT and APP/PS1 cells for background noisy input stimulation at 0.5*Hz* in a realistic, active CA1 model by Poirazi *et al*. (2003b). The inset on the *top right* shows sample trajectories for the voltage response of two sample cells. The mean firing rate is indicated by the dashed lines (*black* WT 25.24 *±* 6.74*Hz*, *orange* APP/PS1 23.69 *±* 3.87*Hz*). **C**, Voltage change versus dendritic length of WT and APP/PS1 cells for three pathway stimulations of 5*Hz* and 0.3 correlation are shown for the passive model by Poirazi *et al*. (2003b). The dashed lines show the mean voltage responses (the asterisk depicts *p <* 0.05). **D**, Firing rates versus dendritic length of WT and APP/PS1 cells for three pathway stimulations of 5*Hz* and 0.3 correlation are shown for the active model by Poirazi *et al*. (2003b). The dashed lines show the mean firing rate (the asterisk depicts *p <* 0.02).

Next, we simulated synaptic activity separately targeting layer-specific subregions of the dendritic tree. In this way, we tested whether a more “clustered” synaptic activation, coming from particular input pathways, leads to a difference in the response behaviour between the two morphology groups. We simulated a rhythmic activation of the synapses (30% synaptic correlation) in each of the three input areas (entorhinal cortex input, CA3 input to oblique and basal dendrites) at theta frequency (5*Hz*), which resembles a prominent behavioural input pattern received by the hippocampus (Bannister and Larkman, 1995; Megías *et al*., 2001; Ang *et al*., 2005; Manns *et al*., 2007; Takahashi and Magee, 2009; López-Madrona *et al*., 2021).

Again, firstly, we looked at the passive cell model exposed to the stimulated pathways (**Figure 2C**). We found similar voltage changes in passive WT and APP/PS1 cells evoked by the perforant pathway stimulation (**Figure 2C**, *Left panel:* WT 20.12 *±* 4.05*mV*, APP/PS1 18.68 *±* 4.72*mV*). However, when stimulating the Schaffer collateral inputs on the apical dendrites, we found slightly larger voltage changes in the APP/PS1 cell group (**Figure 2C**, *Middle panel:* WT 39.48 *±* 7.63*mV*, APP/PS1 43.26 *±* 6.48*mV*; *p <* 0.05). Interestingly, this behaviour was reversed when we looked at the voltage change caused by the CA3 input to the basal dendrites (**Figure 2C**, *Right panel:* WT 28.86 *±* 6.31*mV*, APP/PS1 25.53 *±* 7.10*mV*, *p* = 0.062).

Secondly, we added again active conductances (Poirazi *et al*., 2003b) and observed similar results as the ones shown for the passive model (compare **Figure 2C** and **Figure 2D**). The perforant pathway stimulation led to comparable firing rates in WT and APP/PS1 cells (**Figure 2D**, *Left panel:* WT 54.07 *±* 9.38*Hz*, APP/PS1 53.62 *±* 12.47*Hz*). For the stimulation of apical Schaffer collateral pathway, the APP/PS1 cells displayed slightly but insignificantly larger firing rates when compared to the WT cells (**Figure 2D**, *Middle panel:* WT 84.38*±*14.44*Hz*, APP/PS1 90.31 *±* 13.27*Hz*, *p* = 0.11). In contrast, the CA3 input to the basal dendrites led to a strong correlation between the cell’s firing rate and dendritic length, displaying a significant decrease in the average firing rate in APP/PS1 neurons when compared to WT controls (**Figure 2D**, *Right panel:* WT 82.44 *±* 27.70*Hz*, APP/PS1 67.11 *±* 18.79*Hz*, asterisk depicts *p <* 0.02).

We hypothesised that these pathway-dependent differences in excitability could be due to the variability found in basal/apical dendritic length ratios across the data set (see **Figure 3A**, mean ratio WT 0.55 *±* 0.26, APP/PS1 0.39 *±* 0.22, asterisk depicts *p <* 0.02). A cell with a smaller basal/apical ratio receives relatively less basal input per total cell size (and experiences a larger inactivated apical region that acts as a voltage sink) than a cell with a larger ratio, which could lead to the differences in the firing output.

**Figure 3.**
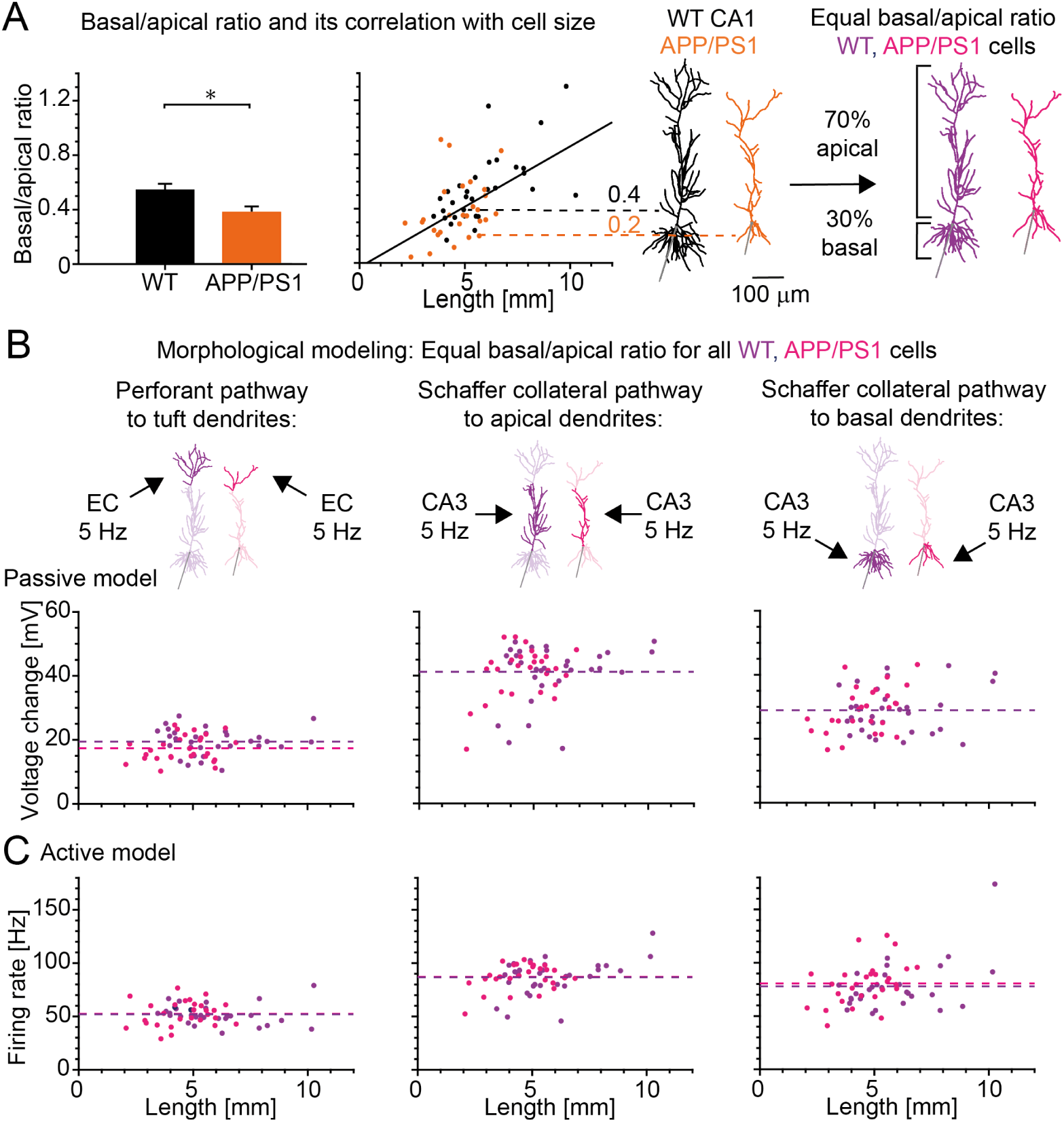
Morphological modelling with artificially equalized basal/apical dendrite ratios reveals similar responses of WT and APP/PS1 model cells to the activation of main input pathways. **A**, *Left*: Comparison of basal/apical ratio between WT and APP/PS1 cells (mean ratio WT in *black*: 0.55 *±* 0.26, APP/PS1 in *orange*: 0.39 *±* 0.22, asterisk depicts *p <* 0.02). *Middle*: Basal/apical ratio versus cell length with the solid black line indicating the linear regression. *Right*: CA1 morphologies remodeled to equal basal/apical ratio of 30%*/*70% for all cells (*dark purple*: WT, *light purple*: APP/PS1). **B & C**, Similar to **Figure 2 C, D** but with scaled morphologies of equal basal/apical ratio.

To test this hypothesis, we used morphological modelling to scale the WT and APP/PS1 cell morphologies to a fixed 30*/*70 basal/apical ratio while keeping the total dendritic length the same as in the original morphologies (**Figure 3A**, *right panel, purple*). We found that, after this structural scaling, the strong correlation between the cell’s output and dendritic length was lost, leading to an almost identical average voltage change and firing rate for both morphology groups (**Figure 3B**, *left*: perforant pathway, WT 18.99 *±* 4.02*mV*, APP/PS1 16.93 *±* 3.91*mV*, *middle*: apical Schaffer collateral, WT 41.15 *±* 8.88*mV*, APP/PS1 41.09 *±* 7.73*mV*, *right*: basal Schaffer collateral, WT 28.88*±*7.25*mV*, APP/PS1 28.92*±*7.10*mV* and **Figure 3C**, *left*: perforant pathway, WT 52.18 *±* 9.34*Hz*, APP/PS1 52.40 *±* 11.79*Hz*, *middle*: apical Schaffer collateral, WT 87.28*±*16.62*Hz*, APP/PS1 87.33*±*12.12*Hz*, *right*: basal Schaffer collateral, WT 78.02*±*23.16*Hz*, APP/PS1 80.10 *±* 20.41*Hz*).

These results indicate that due to stronger degeneration of basal dendrites compared to apical dendrites in APP/PS1 cells the stimulation from CA3 to the oblique dendrites can potentially lead to increased excitability in APP/PS1 cells while the input to the basal dendrites can lead to a decrease in excitability.

### Concomitant network and intrinsic cell changes lead to enhanced excitability in APP/PS1 cells

So far, our modelling has shown that although dendritic degeneration can modulate CA1 PC excitability by selective gating of layer-specific inputs, it cannot fully account for the hyperexcitability observed in APP/PS1 mice. Apart from the morphological changes, which we studied until now, changes in the network input as well as in the intrinsic properties of APP/PS1 CA1 PCs could be responsible for their enhanced firing and bursting. Therefore, our next goal was to use *in silico* simulations to clarify the relative contributions of changes in the network input and ion channels and to predict which changes may be most relevant for AD. Our computational analysis focused on disentangling all the mechanisms for hyperexcitability that have been observed in AD mouse models. It resulted in three potential scenarios with altered extrinsic (network) and intrinsic (cellular) properties.

Importantly, in AD, hyperexcitability has been expressed not only as a general increase in firing rates but also as a change in the firing mode in the form of enhanced burst firing of pyramidal neurons (Chen, 2005; Minkeviciene *et al*., 2009; Kellner *et al*., 2014; Berridge, 2014; Ghatak *et al*., 2019; Müller *et al*., 2021). This shift of the firing mode towards spike bursts has been detected in CA1 pyramidal cells in the APP/PS1 mouse model of AD (Šišková *et al*., 2014). Therefore, we explored, which type of network and/or intrinsic changes can lead to the experimentally observed change in the cell’s firing mode. We identified three scenarios for increased burst firing without changes in solitary spike firing (**Figure 4C-E**) as observed in experimental data (Šišková *et al*., 2014; see their figure 1B).

**Figure 4.**
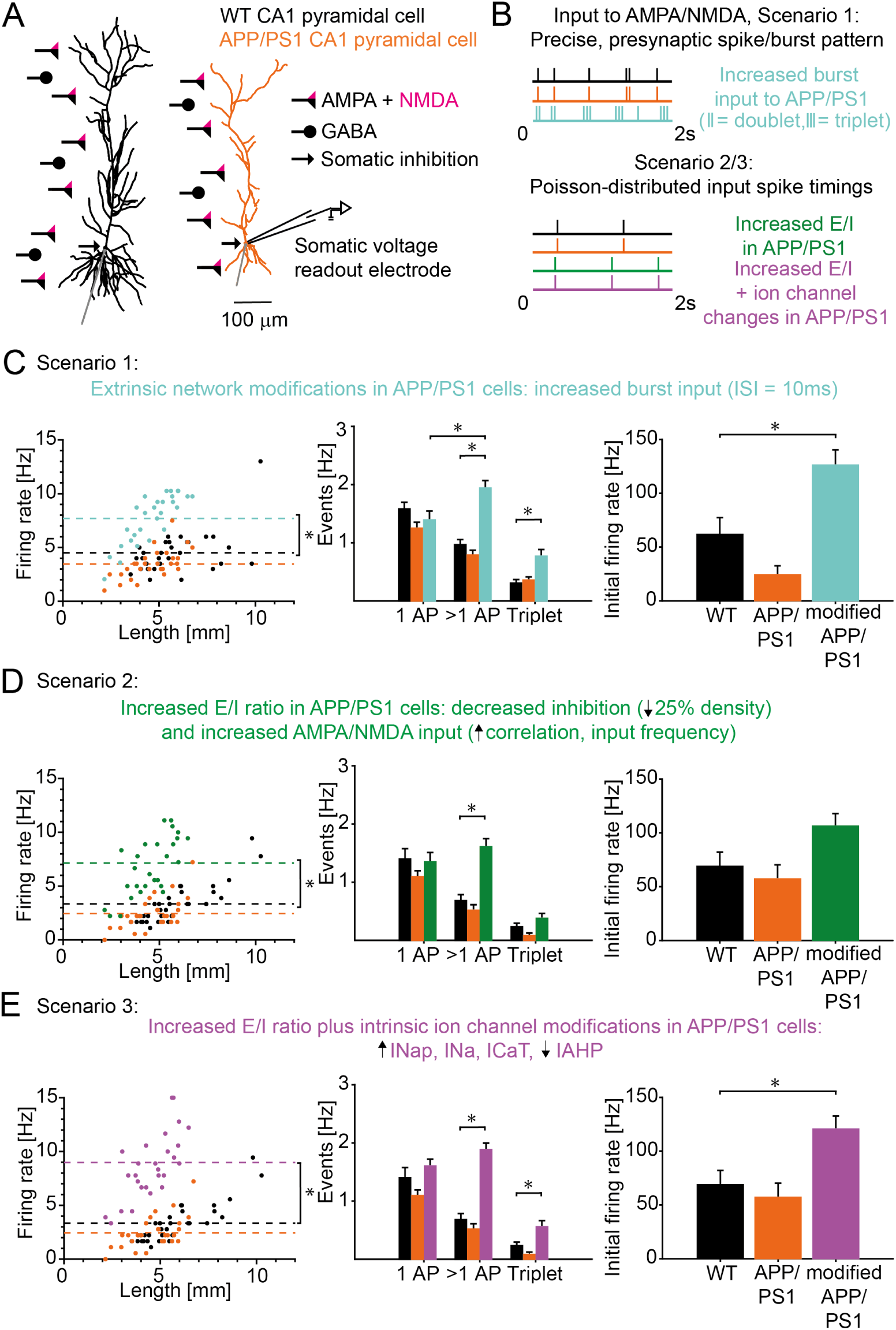
Transition from solitary to burst firing in APP/PS1 cells due to concomitant extrinsic (network) and intrinsic neuronal changes. **A**, Similar organisation as Fig 2A but with GABAergic inhibition extended to somatic region. **B**, Example input pattern to the AMPA/NMDA synapses for Scenario 1 (WT in *black* and APP/PS1 in *orange* receive four singlets and one doublet per two seconds and synapse. Increased burst input to APP/PS1 in *turquoise* with one singlet, two doublets and three triplets per two seconds and synapse) and Scenarios two and three (WT in *black* and APP/PS1 in *orange* with 1*Hz* Poisson input. Increased *E/I* ratio for Scenario 2 in APP/PS1 in *green* with 1.3*Hz* Poisson input. *E/I* ratio increase plus ion channel alterations for Scenario 3 in APP/PS1 in *purple*). **C**, Scenario 1: *Left*, Firing rate versus dendritic length in WT, APP/PS1 and APP/PS1 cells with increased burst input (*turquoise*). The input pattern can be seen in **B**. The dashed lines indicate the mean firing rate. The asterisk depicts *p <* 3 *·* 10*^−^*^8^ for a significant overall firing rate increase. *Middle*, Number of events for the same cell groups: single AP, bursts (*>* 1*AP*, *ISI ≤* 13.3*ms*) and triplets. The asterisk indicates *p <* 5 *·* 10*^−^*^9^ for bursts and *p <* 3 *·* 10*^−^*^4^ for triplets respectively. The mode change from single spikes to predominantly bursts is significant with *p* = 0.008. Note that experiments (Šišková *et al*., 2014) showed unchanged single spikes but strongly increased number of bursts (*>* 1*AP*) and increased triplet AP firing in APP/PS1 pyramidal neurons. *Right*, Initial firing rate between the first two APs (*p <* 2 *·* 10*^−^*^5^). **D**, Scenario 2: *Left*, Firing rate versus dendritic length of WT, APP/PS1 and APP/PS1 cells with increased *E/I* ratio (*green*: 25% decreased somatic and dendritic inhibition, upregulated correlation from 0.4 to 0.8 and Poisson frequency 1*Hz* to 1.3*Hz*, an example input pattern can be seen in **B**). The dashed lines indicate the mean firing rate. The asterisk depicts *p <* 4 *·* 10*^−^*^8^. *Middle*, Number of events for the same cell groups: single AP, bursts and triplets. The asterisk indicates a significant burst increase with *p <* 6 *·* 10*^−^*^7^. *Right*, Initial firing rate of the first two APs. **E**, Scenario 3: *Left*, Firing rate versus dendritic length of WT, APP/PS1 and APP/PS1 cells with the same increase in *E/I* ratio as in Scenario 2 (**D**) but additional, experimentally observed ion channel changes (*purple*: increased *I_Nap_*, *I_Na_*, *I_CaT_* densities, decreased *I_AHP_* density; see details in **Methods**). The dashed lines indicate the mean firing rate. The asterisk depicts a significant increase in firing rate with *p <* 1 *·* 10*^−^*^9^. *Middle*, Number of events: single AP, bursts and triplets. The asterisk indicates significant increase with *p <* 1 *·* 10*^−^*^9^ for bursts and *p <* 0.013 for triplets respectively. *Right*, Initial firing rate of the first two APs (*p <* 0.007).

#### Scenario 1: increased excitatory network burst input

CA1 pyramidal cells are embedded in a wider network with early-onset hyperexcitability (Palop and Mucke, 2016; Zott *et al*., 2018; Selkoe, 2019; Kazim *et al*., 2021). Therefore, we first tested as whether increased network activity with enhanced bursting would transfer to increased burst firing in model neurons. We modeled control network burst input to synapses using four singlets and one doublet per two seconds (’WT’ in **Figure 4C**). In contrast, we modeled enhanced network burst input to synapses using one singlet, two doublets and three triplets per two seconds (’modified APP/PS1’ in **Figure 4C**). The input pattern and time profile can be seen in **Figure 4B** Scenario 1 (coefficient of variation for the input frequency per cell: *cv* = 0, since all stimulated synapses get the same number of input spikes, see **Methods** for simulation details). We observed that these changes in network activity on average led to an increase of the output firing frequency of the APP/PS1 cells (’modified APP/PS1’ in Scenario 1), when compared to the behaviour of both cell groups with a non-hyperexcitable (control) network input (**Figure 4C**, *Left panel*: WT 4.50 *±* 1.98*Hz*, APP/PS1 3.46 *±* 1.36*Hz*, modified APP/PS1 in Scenario 1 7.69 *±* 2.30*Hz* with *p <* 3 *·* 10*^−^*^8^). This overall increase in the cell’s excitability was due to a transition from solitary action potentials (APs) to burst firing of more than one AP (**Figure 4C**, *Middle panel*: WT 0.97 *±* 0.48*Hz*, APP/PS1 0.79 *±* 0.46*Hz*, modified APP/PS1 in Scenario 1 1.94 *±* 0.67*Hz* with *p <* 5 *·* 10*^−^*^9^). Particularly, in line with experimental data in figure 1B of Šišková *et al*. (2014), we observed a significant increase of the number of triplets (**Figure 4C**, *Middle panel*: WT 0.31 *±* 0.33*Hz*, APP/PS1 0.36 *±* 0.30*Hz*, modified APP/PS1 in Scenario 1 0.77*±*0.60*Hz* with *p <* 3*·*10*^−^*^4^), while the single AP firing rate remained constant (**Figure 4C**, *Middle panel*: WT 1.58 *±* 0.65*Hz*, APP/PS1 1.25 *±* 0.55*Hz*, APP/PS1 in Scenario 1 1.39 *±* 0.81*Hz*). This indicates a change of firing mode from predominantly single spikes to burst firing in APP/PS1 cells with increased network activity input in Scenario 1 (*p*_1_*_AP,>_*_1_*_AP_* = 0.008). Additionally, in agreement with data in figure 2B of Šišková *et al*. (2014), the initial firing rate was increased for APP/PS1 cells in Scenario 1 (**Figure 4C**, *Right panel*: WT 54.92 *±* 13.23*Hz*, APP/PS1 24.46 *±* 8.08*Hz*, APP/PS1 in Scenario 1 133.57 *±* 10.34*Hz* with *p <* 2 *·* 10*^−^*^5^).

#### Scenario 2: increased extrinsic ***E/I*** ratio with increased excitatory and decreased inhibitory input

Both enhanced excitatory glutamatergic (Busche and Konnerth, 2016; Zott *et al*., 2019) as well as impaired inhibitory GABAergic transmission have been shown to contribute to neuronal network dysfunction in AD (Busche *et al*., 2008; Schmid *et al*., 2016; Palop and Mucke, 2016; Ambrad Giovannetti and Fuhrmann, 2019; Xu *et al*., 2020; Hijazi *et al*., 2020; Gervais *et al*., 2021). Based on these observations, we tested whether increased *E/I* ratio due to reduced GABAergic inhibition combined with enhanced glutamatergic network input can reproduce experimental data. For this purpose, we increased the *E/I* balance using higher Poisson input frequency (from 1*Hz* to 1.3*Hz*) and stronger spike train correlations (from 0.4 to 0.8) and diminished inhibitory inputs (25% decrease of dendritic and somatic inhibition) in modified APP/PS1 cells of Scenario 2. The input pattern and example time profile can be seen in **Figure 4B** Scenario 2 (coefficient of variation of input per cell, WT: *cv* = 0.41, APP/PS1: *cv* = 0.41 and APP/PS1 of Scenario 2: *cv* = 0.17, this is due to higher mean input frequency and smaller variation, see **Methods** for simulation details). Similarly to Scenario 1, also in Scenario 2 we observed that the overall firing rate (**Figure 4D**, *Left panel*: WT 3.35 *±* 1.87*Hz*, APP/PS1 2.46 *±* 1.48*Hz*, modified APP/PS1 in Scenario 2 7.12 *±* 3.17*Hz* with *p <* 4 *·* 10*^−^*^8^) and especially the number of bursts (**Figure 4D**, *Middle panel*: WT 0.68 *±* 0.59*Hz*, APP/PS1 0.52 *±* 0.52*Hz*, modified APP/PS1 in Scenario 2 1.61 *±* 0.75*Hz* with *p <* 6 *·* 10*^−^*^7^) were significantly increased in the modified APP/PS1 cell group without affecting the single spike rate (**Figure 4D**, *Middle panel*: WT 1.40 *±* 0.99*Hz*, APP/PS1 1.09 *±* 0.55*Hz*, modified APP/PS1 in Scenario 2 1.35 *±* 0.85*Hz*) as reported in figure 1B experiments of Šišková *et al*. (2014). The rate of triplets (**Figure 4D**, *Middle panel*: WT 0.23 *±* 0.34*Hz*, APP/PS1 0.08 *±* 0.25*Hz*, modified APP/PS1 in Scenario 2 0.38 *±* 0.46*Hz* with *p* = 0.28 compared to WT) and the initial firing (**Figure 4D**, *Right panel*: WT 66.65 *±* 13.24*Hz*, APP/PS1 57.82 *±* 13.27*Hz*, modified APP/PS1 in Scenario 2 96.99 *±* 12.58*Hz* with *p* = 0.23 compared to WT) were not significantly different although the trend was similar to data in figure 1B and 2B of Šišková *et al*. (2014).

#### Scenario 3: increased extrinsic *E/I* ratio plus intrinsic ion channel alterations

Ion channel alterations represent another potential mechanism of AD-associated neuronal hyperexcitability. Therefore, in the third scenario, in addition to the enhanced *E/I* ratio, we have included changes in ion channel currents observed in data from literature: downregulation of *I_AHP_*; upregulation of *I_Nap_*, *I_Na_* and *I_CaT_* (see **Methods** for more details) (Yaari *et al*., 2007; Beck and Yaari, 2008; Zhang *et al*., 2014; Wang *et al*., 2015a,b; Liu *et al*., 2015; Wang *et al*., 2016; Ghatak *et al*., 2019; Niday and Bean, 2021; Garg *et al*., 2021). The input pattern and example time profile for Scenario 3 were the same as in Scenario 2 and can be seen in **Figure 4B** (coefficient of variation of input per cell, WT: *cv* = 0.41, APP/PS1: *cv* = 0.41 and modified APP/PS1 in Scenario 3: *cv* = 0.17). Similarly as in Scenario 2, the overall firing rate (**Figure 4E**, *Left panel*:WT 3.35 *±* 1.87*Hz*, APP/PS1 2.46 *±* 1.48*Hz*, modified APP/PS1 in Scenario 3 8.97 *±* 3.51*Hz* with *p <* 1 *·* 10*^−^*^9^) and burst rate were increased (**Figure 4E**, *Middle panel*:WT 0.68 *±* 0.59*Hz*, APP/PS1 0.52 *±* 0.52*Hz*, modified APP/PS1 in Scenario 3 1.88 *±* 0.63*Hz* with *p <* 1 *·* 10*^−^*^9^). At the same time, in line with Šišková *et al*. (2014), the single spike rate stayed the same (**Figure 4E**, *Middle panel*: WT 1.40 *±* 0.99*Hz*, APP/PS1 1.09 *±* 0.55*Hz*, modified APP/PS1 in Scenario 3 1.60 *±* 0.66*Hz*). Moreover, compared to the previous scenario, changes in the intrinsic excitability led to a significant boost in the triplet firing frequency (**Figure 4E**, *Middle panel*: WT 0.23 *±* 0.34*Hz*, APP/PS1 0.08 *±* 0.25*Hz*, modified APP/PS1 in scenario 3 0.56 *±* 0.60*Hz* with *p <* 0.013) and initial firing rate (**Figure 4E**, *Right panel*: WT 66.65 *±* 13.24*Hz*, APP/PS1 57.82 *±* 13.27*Hz*, modified APP/PS1 in scenario 3 118.02 *±* 16.90*Hz* with *p <* 0.007), which improved the quantitative match with electrophysiological data from figure 1B and 2B of Šišková *et al*. (2014). In contrast, increased intrinsic excitability (due to altered ion channel expression) alone did not lead to a transition from solitary to burst firing though the burst firing rate was still significantly increased (**Supplementary Figure S4C**, *Middle panel*: WT 0.48 *±* 0.45*Hz*, APP/PS1 0.33 *±* 0.39*Hz*, modified APP/PS1 with intrinsic ion channel changes 1.15 *±* 0.71*Hz* with *p <* 2 *·* 10*^−^*^5^). The triplet firing in general was enhanced, with a significant increase between original APP/PS1 cells and modified APP/PS1 cells with intrinsic channel changes (**Supplementary Figure S4C**, *Middle panel*: WT 0.17 *±* 0.30*Hz*, APP/PS1 0.041 *±* 0.15*Hz*, modified APP/PS1 with intrinsic ion channel changes 0.22 *±* 0.32*Hz*, *p* = 0.04). When we performed control simulations and applied the same ion channel and network modifications to the WT cells with larger sizes (due to the lack of dendritic degeneration), we found an even higher increase in burst firing compared to the increase in APP/PS1 cells (**Supplementary Figure S5C**, *Right panel*: extrinsic changes in scenario 2 with WT 2.22 *±* 0.85*Hz*, APP/PS1 1.61 *±* 0.75*Hz*, *p* = 0.0164 and extrinsic/intrinsic changes in Scenario 3 with WT 2.46 *±* 0.95*Hz*, APP/PS1 1.88 *±* 0.63*Hz*, *p* = 0.034). This is in agreement with previous studies showing elevated burst firing for larger cells (van Elburg and van Ooyen, 2010). This supports the fact that a smaller dendritic size will reduce the excitability and therefore supports the degeneration as a compensatory mechanism.

When we compare our results of network and ion channel changes, separately and in combination (**Figure 4C-E** and **Supplementary Figure S4C**), we can conclude that the output mode transition from solitary to burst firing that has been observed in APP/PS1 CA1 pyramidal neurons (Šišková *et al*., 2014) can be explained by modified input network dynamics (Scenario 1) or changes in *E/I* balance (Scenario 2) or joint changes in *E/I* balance and ion channels (Scenario 3). The increase in the initial firing rate can be accounted for by Scenarios 1 and 3 or by their combination.

## Discussion

In this study, we explored possible mechanisms accounting for the cellular and network hyperexcitability that has been observed in AD. We investigated the effect of dendritic degeneration, a hallmark of AD, on the cell’s output behaviour. By using detailed compartmental models of CA1 pyramidal neurons based on 3D-reconstructed morphologies from WT and APP/PS1 mice, we showed that the dendritic structural changes in the APP/PS1 morphology group alone cannot explain the increased excitability observed in those cells experimentally. Interestingly, these results suggest that dendritic “atrophy” could actually help maintain the cell’s firing output to distributed synaptic inputs by compensating the loss of synapses through reducing the dendritic arbour and thereby increasing the input resistance. Moreover, when simulating clustered subregion-specific inputs, we observed that stronger degeneration in basal dendrites than apical dendrites leads by itself to decreased responsiveness of APP/PS1 cells to basal synaptic inputs but increased responsiveness to apical inputs. Functional consequences of such selective anatomical gating of layer-specific inputs (due to altered basal/apical dendrite ratio) is unclear. Enhanced responsiveness to apical synaptic activation might partially account for the enhanced firing in APP/PS1 CA1 PCs but cannot explain their enhanced bursting.

Since dendritic degeneration alone could not fully explain hyperexcitability observed in APP/PS1 mice, we consequently investigated potential alternative mechanisms leading to hyperexcitability in the animal model of AD. Our *in silico* analyses identified three scenarios that led to enhanced burst firing: (1) extrinsic changes in the form of increased network burst activity, (2) extrinsic changes in the form of enhanced excitation and reduced inhibition (altered *E/I* ratio) and (3) combined extrinsic and intrinsic changes in the form of increased *E/I* ratio and ion channel modifications. Scenarios 1, 2 and 3 were able to explain not only increased neuronal firing rates but accounted also for the transition of the output mode from solitary to burst firing observed in APP/PS1 CA1 pyramidal neurons (Šišková *et al*., 2014).

### Dendritic degeneration and AD-related hyperexcitability

The basis of cellular hyperactivity in AD remains poorly understood. Several studies have investigated this phenomenon, either by using neuronal cultures and organoids (Ghatak *et al*., 2019), by *in vitro* (Šišková *et al*., 2014) and *in vivo* (Palop *et al*., 2007; Busche *et al*., 2008, 2012; Šišková *et al*., 2014; Verret *et al*., 2012; Busche and Konnerth, 2016; Palop and Mucke, 2016; Sosulina *et al*., 2021) approaches, as well as computational modelling (Šišková *et al*., 2014; van Elburg and van Ooyen, 2010; Vitale *et al*., 2021). Some of these studies suggested that the increased firing and bursting activity of the AD cells could be explained by neurite degeneration (Šišková *et al*., 2014; Ghatak *et al*., 2019) given that reduced dendritic length and complexity promotes the intrinsic excitability and integration of synaptic inputs (van Elburg and van Ooyen, 2010).

These proposals were based on the well known eletrotonic mechanisms (van Ooyen *et al*., 2002) according to which dendritic degeneration reduces neuronal surface and therefore decreases membrane conductance. This makes neurons more excitable in terms of larger voltage responses to somatic current injections as well as to synaptic stimulation Šišková *et al*. (2014). In other words, a small cell with the same number of input synapses as a larger cell shows increased synaptic excitation. However, if also synaptic loss is considered then dendritic shortening with its increase in input resistance may counteract the lower number of synapses (Platschek *et al*., 2016). Although dendritic degeneration decreases the total number of synapses it keeps their relative number (synapse density per surface area) unchanged Šišková *et al*. (2014). In this way, dendritic degeneration could paradoxically help maintain the size invariance of synaptically driven neuronal firing in the presence of rarefied synaptic input. This possibility has been previously shown in electrophysiological and morphological models of dendritic atrophy after entorhinal cortex lesion (Platschek *et al*., 2016). The models have revealed that dendritic atrophy was capable of adjusting the excitability of neurons thus compensating for the denervation-evoked loss of synapses (Platschek *et al*., 2016, 2017). This computational principle has recently been generalised to all cell types with a variety of dendritic shapes and sizes (Cuntz *et al*., 2021). Mathematical analysis and numerical simulations have shown that neuronal excitation in response to distributed inputs was largely unaffected by the dendrite length if synaptic density was kept constant (Cuntz *et al*., 2021).

Similarly, our detailed computational modelling here indicates that atrophied APP/PS1 cell morphologies probably do not lead on their own to increased firing and bursting as compared to the WT group, when driven by the same density of distributed synaptic inputs (**Figure 1** and **Figure 2**). The condition of same synaptic density is based on similar spine density in APP/PS1 and WT dendrites (Šišková *et al*., 2014). Later stages of the AD progression (*>* 20 months) show further spine loss and a decrease in spine density (Spires-Jones *et al*., 2007; Knobloch and Mansuy, 2008; Bittner *et al*., 2010).

Our results regarding excitability and dendritic degeneration remained consistent for both simple synaptic models with AMPA synapses (**Figure 1**) as well as for more realistic and detailed synaptic models with distributed AMPA, NMDA and GABA-A synapses (**Figure 2B**). This was not only true for changes in dendritic length in the APP/PS1 cell morphologies but also for other morphological measures such as the number of branch-points or the surface area of the cells. In each case we found no hyperactivity in the APP/PS1 cells, which display dendritic degeneration (**Supplementary Figure S2**). To test the robustness of our findings with respect to biophysical parameters, we implemented a second compartmental model by Jarsky *et al*. (2005), which confirmed the results of the model by Poirazi *et al*. (2003b) by showing no hyperexcitability in APP/PS1 cells due to purely morphological changes (Supplementary Figure S3).

We further investigated the effect of specific pathway inputs on the cell’s response for both morphology groups and found a link between dendritic degeneration and change in excitability (**Figure 2**). Stimulation of basal dendrites resulted in decreased firing of APP/PS1 cells compared to WT cells. Conversely, stimulation of apical dendrites increased the activity of APP/PS1 cells though it was only significant in the passive model (**Figure 2C, D**, *Middle panels*). This phenomenon is explained by a decrease in basal/apical ratio in APP/PS1 cells due to stronger basal degeneration resulting in less stimulated area per whole cell surface when only basal dendrites were activated. Vice versa, the stimulated area per whole cell surface was increased when only apical dendrites were activated (**Figure 3A**, *Left*). The different proportions of basal and apical dendrites therefore led to a selective gating of inputs from CA3. Interestingly, in case of equal basal/apical ratio this difference in excitation disappeared as shown by the morphological modelling of artificially scaled WT and APP/PS1 cells in **Figure 3**.

Taken together, our modelling indicates that additional, non-morphological changes are required to trigger increased burst firing in CA1 pyramidal neurons of APP/PS1 mice as reported in *in vivo* whole-cell and LFP recordings (Šišková *et al*. 2014; see their figure 1B). Our simulation results (**Figure 1, 2** and **Supplementary Figure S3**) show that the APP/PS1 cells display no hyperactivity compared to the WT cells even though they are on average smaller (**Supplementary Figure S2B**) and have less synapses. We therefore propose that dendritic atrophy might be a compensatory mechanism contributing to firing rate homeostasis (Platschek *et al*., 2016, 2017). The observation of morphological compensatory effects counteracting the functional impairments observed in AD have been repeatedly reported. Modifications such as an increase in size of remaining dendritic spines after AD-related spine loss (Fiala *et al*., 2002; Dickstein *et al*., 2010; Neuman *et al*., 2015) and changes in the topology and size of dendrites (Graveland *et al*., 1985; Arendt *et al*., 1995) seem to point towards the stabilising effect of spine and dendritic remodelling. Dendritic degeneration not only changes the overall size and complexity of the cell, but also the amount of membrane and available space for synaptic contacts. In fact, saving energy resources by reducing available surface membrane after synapse loss appears to be one of the reasons for dendritic retraction. Accordingly, dendritic retraction has been observed as a consequence of presynaptic neuron loss and an associated presynaptic denervation and synapse loss in AD but also other neuronal lesions (Platschek *et al*., 2017). However, the neuronal output is not necessarily decreased because the reduction in the number of synapses is well compensated morphologically by an increased input resistance (Platschek *et al*., 2016; Cuntz *et al*., 2021). In this way, dendritic changes that occur after the synapse loss observed in AD, might help restore the input-output function of the cell. Such input-output homeostasis would be beneficial not only for the single neuron, but also for the network in which it is embedded. Similar compensatory strategies have been reported across species (Cuntz *et al*., 2013) and cell types (Weaver and Wearne, 2008; Tripodi *et al*., 2008; Platschek *et al*., 2016). In line with this idea, other studies have reported a conservation of the electrophysiological properties despite structural changes of AD-affected neurons (Rocher *et al*., 2008; Somogyi *et al*., 2016). Moreover, although it is commonly assumed that cells possess a specific structure in order to support a specific function, it has been recently shown that dendrite morphology can be well predicted by anatomical connectivity (Cuntz *et al*., 2010) and that the cell’s size and shape is largely independent of it’s output under controlled conditions for distributed synaptic activation (Cuntz *et al*., 2021). This suggests that early AD-related modifications in the cellular structure (dendritic atrophy) compensate for synaptic loss as long as possible and keep the function intact.

### Alterations in extrinsic *E/I* balance and intrinsic ion channel properties

Since neuronal hyperexcitability in the APP/PS1 cells *in vivo* does not seem to be mediated by isolated changes in the dendritic morphology, we explored possible alternative explanations for its origin. Previous studies have observed an increased network activity in AD-affected brain regions (Šišková *et al*., 2014; Palop and Mucke, 2016; Zott *et al*., 2018; Selkoe, 2019; Maestú *et al*., 2021; Kazim *et al*., 2021), which eventually can lead to epileptiform activity (Minkeviciene *et al*., 2009; Noebels, 2011). We found that increasing the excitability of the network by enhancing the excitatory input that feeds to our APP/PS1 CA1 pyramidal model cells can sufficiently boost their bursting frequency and alter the predominant mode of firing from single spike to burst firing (**Figure 4C**) as observed in APP/PS1 mice (Šišková *et al*., 2014; see their figure 1B). Thus, our modelling predicts a significant contribution of extrinsic network properties to network hyperactivation. Since neurons are embedded in a network of other AD-affected neurons, this could lead to a cascade effect, amplifying the network hyperactivity.

The hyperexcitability in AD cells has also been previously linked to ion channel modifications (Chen, 2005; Beck and Yaari, 2008; Brown *et al*., 2011; Zhang *et al*., 2014; Kerrigan *et al*., 2014; Liu *et al*., 2015; Wang *et al*., 2015b,a, 2016; Musial *et al*., 2018; Ghatak *et al*., 2019; Villa *et al*., 2020; Vitale *et al*., 2021; Müller *et al*., 2021). Therefore, we explored which combination of these previously reported ion channel changes led to an increase of excitability in the APP/PS1 morphologies. In line with experimental findings, we found several ion channel changes that increased the overall spike rate and burst firing: down-regulation of *I_AHP_* (Beck and Yaari, 2008; Zhang *et al*., 2014; Wang *et al*., 2015a; Niday and Bean, 2021), up-regulation of *I_Na_* and persistent *I_Nap_* (Williams and Stuart, 1999; Yue *et al*., 2005; Beck and Yaari, 2008; Liu *et al*., 2015; Wang *et al*., 2016; Ghatak *et al*., 2019) and up-regulation of T-type calcium current *I_CaT_* (Yaari *et al*., 2007; Beck and Yaari, 2008; Cain and Snutch, 2013; Medlock *et al*., 2018; Garg *et al*., 2021). Our simulations showed that, although ion channel modifications alone led to an increased firing of especially bursts (**Supplementary Figure S4C**, *Middle panel*), they did not reproduce quantitatively the output mode transition from single spiking to predominantly burst firing as reported in figure 1B of (Šišková *et al*., 2014). We tested also changes of other ion channels that have been linked to hyperexcitability including potassium, L-type calcium and hyperpolarisation-activated, cyclic nucleotide-gated *HCN* channels (Beck and Yaari, 2008; Musial *et al*., 2018; Vitale *et al*., 2021) but our simulations did not show a significant contribution to the increase in burst activity (see Methods).

However, importantly, the combination of these intrinsic ion channel modifications together with extrinsic network changes in the form of increased input frequency and input correlations in excitatory connections (AMPA/NMDA) and decreased inhibitory synaptic activity (25% reduction) led not only to elevated firing rates but also to an output mode switch from solitary to burst firing in the APP/PS1 morphology group (**Figure 4E**). Such changes in *E/I* balance as a result of reduced inhibition and increased excitation have been shown previously in AD mouse models (Busche *et al*., 2008; Schmid *et al*., 2016; Palop *et al*., 2007; Palop and Mucke, 2016; Ambrad Giovannetti and Fuhrmann, 2019; Xu *et al*., 2020; Hijazi *et al*., 2020; Gervais *et al*., 2021). Our results show that joint alterations in *E/I* balance and ion channels can explain not only qualitatively but also quantitatively the AD-related hyperexcitability including the shift from single spikes to bursts of spikes. Interestingly, when we applied the same ion channel and network modifications to the WT morphologies we found an even higher increase in burst firing compared to the APP/PS1 cells (**Supplementary Figure S5C** and compare *left* and *middle* for WT and APP/PS1 cells respectively). A plausible explanation for this is that the observed dendritic degeneration can partially counter the increase in burst firing in APP/PS1 cells. This is in agreement with previous computational and morphological studies showing the effect of dendritic size and topology on the bursting behaviour of pyramidal cells (van Elburg and van Ooyen, 2010).

In this study, we focused on stages of AD progression, at which amyloid plaques can be reliably observed. A recent study in transgenic rats has provided *in vivo* evidence that, at an early pre-plaque stage, CA1 hyperexcitability is mediated predominantly by increased intrinsic excitability (Sosulina *et al*., 2021). Accordingly, the dysfunction of inhibitory and excitatory transmission (Busche and Konnerth, 2016) would be expected to play a role at later stages of AD although the precise sequence of pathophysiological events remains to be established. It would be insightful to model mechanistically earlier stages and compare them to the later stages capturing not only the later-stage synergy of extrinsic and intrinsic pathomechanisms but also their initial sequence and dynamics in time. First steps towards such stage-dependent computational synthesis have been recently reported in a study of APP/PS1 mice at three different ages (Vitale *et al*., 2021). This work revealed age-dependent changes in membrane time constant, expression of HCN channels, AP width and firing behaviour (evoked by somatic current injections) in APP/PS1 animals. Although measuring and modelling firing behaviour triggered by somatic current injections is important, our study shows that for a better understanding of age-depedendent progress in AD-related hyperexcitability of single cells and networks, it will be important to characterise and model also the more natural, i.e. synaptically driven, firing behaviour.

### Limitations and applicability

Ion channel degeneracy (Goaillard and Marder, 2021) implies that neurons of the same cell type can incorporate different densities of multiple ion channel types that have overlapping properties and contribute to the same neuronal response (Drion *et al*., 2015; O’Leary, 2018; Schneider *et al*., 2021). Therefore, with respect to ion channel degeneracy and variability it is likely that individual AD-affected CA1 pyramidal cells react differently with a variety of ion channel modifications which nevertheless can lead to the same neuronal hyperactivity with the increased firing rates and the burst rates. In addition, depending on the stage of the disease, different proportion of neurons may be affected by AD with a subset of them displaying hyperexcitability or hypoexcitability (Busche *et al*., 2008; Zott *et al*., 2018; Dunn and Kaczorowski, 2019) further increasing the variability in the network. In a healthy brain, the variability in ion channel expression is normally a compensatory homeostatic mechanism. However, the compensation might only be possible for a limited range of parameters or fail due to the complex interactions in the high-dimensional parameter space of the extrinsic and intrinsic neuronal mechanisms (O’Leary, 2018). Indeed, the failure of homeostatic machinery including dysregulation of firing rate homeostasis is one of the main hypotheses for the vicious circle of AD progression (Frere and Slutsky, 2018; Dunn and Kaczorowski, 2019). Future modelling and experimental studies should address the variability in AD-related ion channel changes and in the failure of firing homeostasis.

Another limitation of our study is that there are additional mechanisms underlying AD-associated changes in extrinsic and intrinsic properties that we have not systematically tested in our simulations. For example, amyloid-beta accumulation as one of the hallmarks of AD progression may affect extracellular glutamate concentrations (Bezprozvanny and Mattson, 2008; Scimemi *et al*., 2013; Bao *et al*., 2021). Our simulations showed that indeed an increased NMDA time constant, which represented a delayed clearance of synaptically released glutamate, can result in an increased firing rate of APP/PS1 cells (**Supplementary Figure S4D**). However, the burst firing rate remained unchanged with only the single AP firing rate being elevated indicating that other changes are necessary to fully explain hyperexcitability data. The impairment of synaptic plasticity due to amyloid-beta accumulation is a further indicator of AD leading to inhibited LTP and memory deficits (Rowan *et al*., 2003; Shankar *et al*., 2008). This impairment however may rather counteract the hyperexcitability observed in the APP/PS1 morphologies. Axon initial segment (AIS) properties are well known as an important factor for neuronal excitability (Kuba *et al*., 2014; Evans *et al*., 2015). Impaired AIS plasticity due to AD-related Amyloid-beta proximity can therefore lead to alterations in the cellular excitability (León-Espinosa *et al*., 2012; Sun *et al*., 2014; Zhang *et al*., 2014; Dongmin Sohn *et al*., 2019; Booker *et al*., 2020). We did not include AIS changes in our study since we have no anatomical information for the examined morphologies. However, future work should investigate the role of the AIS and its potential changes in AD-related hyperactivity.

An additional caveat is that our work focused specifically on the AD-related hyperexcitability in the hippocampus of APP/PS1 mice. Hyperexcitability in other brain areas in these and other AD mice might be a result of other or additional circuit mechanisms. Indeed, a recent study has shown that early AD-related hyperexcitability in the somatosensory cortex of the familial AD mouse model is a consequence of a dysfunction in the firing of GABAergic parvalbumin interneurons due to alterations of their potassium (Kv3) channels (Olah *et al*., 2022).

### Degeneracy in AD-related hyperexcitability

Our findings are in agreement with a general and increasingly accepted concept of degeneracy (Edelman and Gally, 2001). In complex degenerate systems, such as the nervous system, multiple distinct mechanisms can be sufficient, but typically not necessary, to account for a given function or malfunction, such as normal excitability or hyperexcitability (Kamaleddin, 2022; see also Neymotin *et al*., 2016; Ratté and Prescott, 2016; O’Leary, 2018; Medlock *et al*., 2022). Consistent with the degeneracy framework, our results suggest that multiple extrinsic and intrinsic mechanisms are sufficient but not necessary to increase neuronal excitability in AD. More specifically, with respect to the altered spike pattern in AD mice (Šišková *et al*., 2014), we show that enhanced network burst input (Scenario 1), enhanced *E/I* ratio (Scenario 2) or enhanced *E/I* ratio together with ion channel changes (Scenario 3) are sufficient to account for the transition of output mode from solitary to burst firing.

Paradoxically, in systems displaying degeneracy, repairing a single target mechanism may not be enough for restoring their normal excitability (Kamaleddin, 2022). Compensatory effects and adaptations (O’Leary, 2018) and their variability in different individuals (Sakurai *et al*., 2014; Onasch and Gjorgjieva, 2020; Medlock *et al*., 2022) can sometimes impede mono-causal therapeutic options targeting a single mechanism (Ratté *et al*., 2014; Ratté and Prescott, 2016). Therefore, the framework of degeneracy may help develop multi-causal intervention strategies, which could include multiple sets of extrinsic and intrinsic perturbations rescuing pathologically increased excitability in AD.

## Author contributions

M.M., L.M., S.R., H.C., and P.J. designed the study. M.M. and L.M. performed the simulations and analysed the data. M.M., L.M., S.R., H.C., and P.J. wrote the paper.

## Material and methods

### Data analysis and code availability

Morphological analysis was performed in Matlab (Mathworks Inc, version 2018a) using our own software package, the TREES toolbox (Cuntz *et al*. (2011), www.treestoolbox.org). All passive and active compartmental model simulations were run in NEURON (Hines and Carnevale, 2004) via a newly developed software to control NEURON with Matlab and the TREES toolbox, the TREES-to-NEURON (T2N) interface (Beining *et al*., 2017). The results and figure panels throughout the manuscript were further analysed and generated with Matlab and Adobe Illustrator CS6.

The code will be made available online.

### Dendrite morphologies

The morphology data set used in the study included 59 mouse hippocampal CA1 pyramidal cell reconstructions from 10-14 months old WT (*n* = 31) and APP/PS1 (*n* = 28) mice (Šišková *et al*., 2014). Each 3D reconstruction was translated to a format supported by Matlab and carefully examined for any morphological inconsistencies and physical integrity (see distribution of cells by length and two example morphologies in **Supplementary Figure S2A**, *left*). Preprocessing of the morphologies included therefore the removal of unrealistic nodes and trifurcations (*`repair tree`*, *`clean tree`*) and the correction of unrealistic sudden shifts especially in the *z*-axis (*`zcorr tree`*, *`smooth tree`*). When divided, the separated basal and apical subtrees had to be concatenated. All nodes from the reconstruction were redistributed to have equal inter-nodal distances of 1*µm* for a high structural resolution (*`resample tree`*). The dendritic diameter was tapered (quadratic taper: *`quadfit tree`*, *`quaddiameter tree`* as in Cuntz *et al*. (2007)) in order to correct for unrealistically distributed diameters in the original reconstructions. Statistical comparison of average dendritic path length (WT 5.19 *±* 1.75*mm*, APP/PS1 4.00 *±* 1.22*mm*, *p* = 0.004), number of branching points (WT 52.16 *±* 17.79, APP/PS1 41.61 *±* 11.67, *p* = 0.01) and dendrite surface area (WT 0.026 *±* 0.008*mm*^2^, APP/PS1 0.02 *±* 0.008*mm*^2^, *p* = 0.007) between WT and APP/PS1 model cell groups showed significant differences (**Supplementary Figure S2A**, *right*). The average dendritic diameter was similar in both cell groups (WT 1.6 *±* 0.24*µm*, APP/PS1 1.57 *±* 0.23*µm*, *p* = 0.6). However, pyramidal cells typically depict an average dendritic diameter of 0.8*µm* (Benavides-Piccione *et al*., 2020). Finally, the dendrites were therefore normalised to an average diameter of 1*µm* as part of the morphologies preprocessing as described previously (Cuntz *et al*., 2021). Control simulations with diameter 0.8*µm* showed similar results and can be seen in **Supplementary Figure S6**. None of the reconstructions included a soma nor an axon. Therefore, an artificial cosine shaped soma was added having a surface area of 560*µm*^2^ (*≈* 4 *·* 137*µm*^2^, which is the soma perimeter) and a maximum diameter of 10*µm*, corresponding to the typical average size of somata in pyramidal neurons (Benavides-Piccione *et al*., 2020). The same axon was appended for consistency to the soma of all neurons. The axon was created using several concatenated cylindrical compartments, presenting a total length of 630*µm* and an average diameter of 0.5*µm*, modelling the axon hillock, initial segment, five nodes of Ranvier and myelination (Benavides-Piccione *et al*., 2020). For the simulations in **Figure 3** the original WT and APP/PS1 cell morphologies were artificially scaled in order to obtain specific basal/apical length ratios of 30*/*70. Each cell morphology kept its original total dendritic length, varying only the relative length proportion between the basal and the apical dendrites.

### Normalisation of laminar structure and dendritic regions

In order to evaluate the influence of morphology alone on the cell’s output, the biophysical properties with electrotonic parameters and active mechanisms must be consistently distributed. Therefore, a comprehensive way to define subregions with distance boundaries was needed. To do this, we defined a normalised division of the CA1 dendrites into the different hippocampal layers and dendritic subregions. The anatomical boundaries of the CA1 hippocampal area were delimited depending on the contribution of the dendritic length to each layer (Bannister and Larkman, 1995; Trommald *et al*., 1995; Megías *et al*., 2001), obtaining that, on average, the stratum oriens, stratum radiatum and stratum lacunosum-moleculare represented a 40%, 40% and 20% of the total CA1 dendritic length, respectively. Two major division planes were defined, the SO/SP plane that cut through the cell soma and was orthogonal to the direction of growth of the apical dendrite; and the SR/SLM boundary, which was parallel to the SO/SP layer, leaving in between both planes the relative contribution (67%) of the SR to the apical dendritic length (**Supplementary Figure S1A**). The CA1 pyramidal cell structure was further divided into dendritic sublayers, which were used as inflexion points for changes in electrotonic properties and active channels along the somato-dendritic axis. Four distance inflexion points appear in the majority of CA1 cell models available in *ModelDB*: 100*µm,* 300*µm,* 350*µm* and 500*µm* path length (or laminar depth). Each of these distance values was translated into dendritic apical length contribution by calculating the amount of dendritic length that lies within the different distance limits for both ion channel distribution approaches, path and perpendicular length, and expressed as percentage of apical length, using a large data set of CA1 rat cell morphologies available at *NeuroMorpho.Org*. The dendritic apical length within each of the boundaries did not differ substantially, estimating a contribution of 8%, 48%, 67% (matches the SR-SLM boundary) and 85% of total apical length for each distance limit respectively. The entire path from each terminal branch tip that invaded the SLM was drawn. Those nodes, shared by more than half of the paths which were below the SR-SLM boundary, were defined as the main apical dendrite (trunk). The nodes above the SR-SLM limit were designated as the apical tuft. The branches stemming from the trunk that failed to invade the SLM (less than 2*/*3 of the total branch length trespassed the SLM) were labelled as oblique dendrites.

### Biophysical properties of CA1 cell models

The determination of normalised dendritic regions in morphologies of different shapes and sizes (as described above) allowed us to design a coherent morphology-independent way of generalising existing biophysical CA1 pyramidal cell (PC) models to diverse morphologies. For the results in **Figures 1, 2, 3, 4** and **Supplementary Figures S4, S5**, we implemented a biophysically realistic model based on the model developed and validated by Poirazi *et al*. (2003b,a), see model #20212 in ModelDB (Hines and Carnevale, 2004; McDougal *et al*., 2017) translated to T2N in Cuntz *et al*. (2021). The model consisted of active and passive membrane mechanisms including 16 types of ion channels, most of them non-uniformly distributed along the somato-dendritic axis (**Supplementary Figure S1C**). NEURON channel models were used with no modification. The number of segments per section was defined each 75*µm*.

In order to implement the same non-uniform distribution of channel densities for different pyramidal cells whose apical trunk size can vary greatly (as well as its overall size), the original morphology’s trunk laminar depth in the model by Poirazi *et al*. (2003b) was extracted and adjusted to fit the trunk size from each of the cells in the data set. This means, a vector from 0 to a maximum distance of 423.75*µm* (maximum laminar depth of the original cell) was generated, and rescaled at different intervals that depended on the size of each new cell’s apical trunk, but that kept the same minimum and maximum depth values (**Supplementary Figure S1B**). As the channel densities were multiplied by these distance values, every cell had the same minimum and maximum conductance value for the different channels, and kept a proportional distance dependency of conductance changes (same distribution function), when compared to the original model cell.

The dendritic region specification employed for the channel distribution in this model was based on the archetypical regions in pyramidal cells, where the cells are divided into basal, somatic, trunk, apical (oblique and tuft) and axonal regions as described in detail above and shown in **Supplementary Figure S1A**. In addition, the membrane properties for the oblique side branches within the first 50*µm* from the trunk were set to follow the respective trunk conductance values. This extra dendritic region was defined as the peritrunk area. Likewise, distal apical regions, defined by the distal SR and tuft layer boundaries, also possess modified conductance densities in order to account for some specific distal changes in channel distribution. The model does not include any compensation for spines.

For the **Supplementary Figure S3** as a comparison to the model by Poirazi *et al*. (2003b) we also implemented the model by Jarsky *et al*. (2005), see model #116084 in ModelDB. This model included uniform passive parameters throughout the cell and four different types of active conductances: a voltage-gated sodium conductance, a delayed-rectifier potassium conductance, and a proximal and distal A-type potassium conductance as seen in **Figure S3A**. All NEURON channel models were used with no modification. The dendritic region specification employed for the channel distribution in this model was based on the dendritic sublayer division as in **Supplementary Figure S1A**, where the cells were divided into basal, somatic, proximal SR, middle SR, distal SR, tuft (proximal SLM and distal SLM) and axonal regions. Moreover, this model included a spine-correction mechanism that multiplies the specific membrane capacitance and divides the specific membrane resistance for regions above the distal SR by two. For both models the passive and spiking properties were evaluated in **Supplementary Tables S1 and S2**.

### Excitatory and inhibitory synapses

AMPA synapses were used as excitatory input for both active CA1 pyramidal cell spiking models by Jarsky *et al*. (2005) and Poirazi *et al*. (2003b). Synapses were implemented as a dual-exponential of time-dependent conductances with rise time constant of *τ*_1_ = 0.2*ms* and decay time constant of *τ*_2_ = 2.5*ms* while having a reversal potential of 0*mV* (*`Exp2Syn`* NEURON object). The synapses were driven by *`VecStim`* point processes in artificial point neurons with generated Poisson spike trains (T2N function *`t2n poissonSpikeGen`*). The specific Poisson process frequency varied per protocol and is indicated for each simulation in the figure caption. The synaptic weights had a strength of 0.1*nS* and synapse density was homogeneous with 0.5 *· synapses/µm* for all simulations in **Figures 1, S2 and S3** in which only AMPA was implemented.

For the **Figures 2, 3, 4, S4, S5 and S6** additionally to AMPA synapses we implemented NMDA synapses with slower kinetics and nonlinear voltage-dependence. The NMDA synapses included a magnesium block removed by sufficient depolarisation. The synapses followed a dual-exponential slower than the AMPA synapses with rise time constant of *τ*_1_ = 0.33*ms* and decay time constant of *τ*_2_ = 50*ms* while having a reversal potential of 0*mV* (*`Exp2nmda2`* NEURON object based on the model from Krueppel *et al*. (2011)). For the simulation of the Amyloid-beta related prolonged extracellular glutamate transients in **Supplementary Figure S4D** we increased *τ*_2_ to 100*ms*. All NMDA synapses had the same location as the AMPA synapses and were driven by the same presynaptic Poisson processes.

To model inhibitory synapses, we included GABA-A synapses to the spiking model by Poirazi *et al*. (2003b). GABA-A synapses were implemented by using the *`Exp2Syn`* NEURON object with the same kinetics as for the AMPA synapses but with a changed reversal potential of *−*70*mV* . The weights of the GABA-A synapses had a strength of 2*nS* corresponding to values found by Bloss *et al*. (2016).

For more realistic simulations of synaptic distribution, the densities of the AMPA, NMDA and GABA-A synapses were changed depending on the dendritic region, based on the data from Megías *et al*. (2001); Šišková *et al*. (2014); Bloss *et al*. (2016) (**Supplementary Table S3**). It has been shown that apical synaptic weights in CA1 pyramidal cells are not constant but increase for AMPA (decrease for NMDA) synapses with distance to soma (Magee and Cook, 2000; Katz *et al*., 2009; Kim *et al*., 2015). Therefore, to implement a realistic inhomogeneous synaptic weight distribution, the apical AMPA synapses were upscaled and NMDA downscaled in a linear distance-dependent manner for the model by Poirazi *et al*. (2003b). Along the somato-dendritic axis the AMPA synapses followed the relation *g_AMP A_* = (0.4 + *p*) *· scale_AMP A_* for the conductance values, while *scale_AMP A_* corresponded to 0.1*nS* (compare the previous, constant AMPA conductance value of 0.1*nS* in **Figure 1**) and *p* corresponded to the pathlength from soma to synapse location *x* normalised to the maximum path-length of the cell *p* = *path*(*x*)*/max*(*path*(*cell*)) resulting in increasing weights with increasing distance to soma with a strength of 0.04*nS* to 0.14*nS*. The NMDA synapses followed the relation *g_NMDA_*= (1.2 *−* 0.4 *· p*) *· scale_NMDA_* with *scale_NMDA_* = 0.1*nS* resulting in decreasing NMDA weights with increasing distance to soma with a strength of 0.08*nS* to 0.12*nS*. The slope was estimated by applying depolarising inputs to the morphology trunks and adjusting the weights depending on distance to soma in order to guarantee the same somatic EPSP in agreement with data showing dendritic democracy (Magee and Cook, 2000; Häusser, 2001). In line with experimental data (Bittner *et al*., 2012), synaptic weights in the tuft stayed constant with the maximum distance value of the aforementioned relations. At the same time, a lognormal (*µ* = *−*1.5*, σ* = 0.9) term was added to the excitatory conductances (weighted with *scale_AMP A_/*2 and *scale_NMDA_/*2 respectively) to increase their variance and convert the weights into a more realistic distribution with a majority of weak and a minority of strong weights (Ballesteros-Yáñez *et al*., 2006; Arellano *et al*., 2007; Katz *et al*., 2009; Benavides-Piccione *et al*., 2013; Bromer *et al*., 2018) (**Figure 2A**, AMPA inset).

### Synaptic activation of model CA1 pyramidal cells

The synaptic inputs were modelled as transient conductance changes (see above) following APs generated from a presynaptic spike generator. In all simulations the CA1 model cells were subject to ongoing background activity generated randomly as poisson spike trains, with an average spiking frequency that ranged from 0.1*Hz* to 10*Hz* for AMPA synapses in passive and active models with biophysics by Poirazi *et al*. (2003b) in **Figure 1** and 0.5*Hz* to 15*Hz* in models with biophysics by Jarsky *et al*. (2005) in **Supplementary Figure S3**. The AMPA, NMDA and GABA synapses in active spiking models in **Figure 2** and **Figure 3** received an input of 0.5*Hz* for both, WT and APP/PS1 group. We increased the excitatory input frequency to AMPA/NMDA inputs in **Figure 4** and the **Supplementary Figures S4, S5 and S6** to 1*Hz*. In combination with strong perisomatic inhibition in the high *γ* range of 50 *−* 100*Hz* (Craig and McBain, 2015; Strüber *et al*., 2017), the stimulation was sufficient to create spontaneous burst firing patterns (inter-spike-interval *ISI ≤* 13.3*ms*).

To mimic enhanced bursting input from presynaptic AD-affected neuronal partners, we fed spike trains of elevated burst activity to 4% of the excitatory synapses of the CA1 PCs (proportional to the synapse density in CA1 PCs **Supplementary Table S3**) as seen in Scenario 1 in **Figure 4C** in order to generate bursting behaviour similar to experiments by Šišková *et al*. (2014). According to the data range of the overall firing rate and burst pattern in healthy cells of patch-clamp recordings in Šišková *et al*. (2014), WT and APP/PS1 cells received four singlets and one doublet per two seconds per synapse (majority of single spikes, input frequency 3*Hz*, input burst frequency 0.5*Hz*, *ISI* = 10*ms*) whereas the “extrinsic” APP/PS1 cells of Scenario 1 received only one singlet but elevated bursting with two doublets and three triplets per two seconds (majority of bursts, input frequency 7*Hz*, input burst frequency 2.5*Hz*) as seen in the example input pattern in **Figure 4B** and in the overview of the three scenarios in **Supplementary Table S4**. To test the effects of an increased *E/I* ratio the “extrinsic” APP/PS1 cell group of Scenario 2 and the “extrinsic/intrinsic” APP/PS1 cell group of Scenario 3 in **Figure 4D, E** had a higher Poisson input frequency of 1.3*Hz*. All specific input frequencies used in the simulations are indicated in their respective figures. In addition to spontaneous background activity, neuronal firing was driven also by specific input pathways from its connecting network, which was implicitly simulated using spike generators. Therefore, on top of background noise we added correlated theta input of 5*Hz* to the excitatory synapses in tuft (perforant pathway), basal (Schaffer collateral basal pathway) and apical (Schaffer collateral apical pathway) dendritic regions of the CA1 cells (Bannister and Larkman, 1995; Megías *et al*., 2001; Ang *et al*., 2005; Manns *et al*., 2007; Takahashi and Magee, 2009; López-Madrona *et al*., 2021), respectively in **Figure 2** and **Figure 3**. Thereby, to guarantee a correlation of 0.3 of the input spike trains from each pathway, a modified spike generator was used that kept a random spike train with a probability equal to the target correlation for the next synapse’s input spike train (*`t2n poissonSpikeGen2`*). In Scenarios 2 and 3 the input correlation of the Poisson spike trains was increased from 0.4 in the WT and APP/PS1 cell groups to 0.8 in the “extrinsic” and “extrinsic/intrinsic” APP/PS1 cell groups for simulations in **Figure 4D, E** and **Supplementary Figures S5 and S6**. Additional effects of a changed *E/I* network balance due to reduced inhibitory drive from interneurons were modeled by decreasing the density of activated inhibitory synapses in somatic and dendritic regions by 25% proportional to the synapse density in **Supplementary Table S3** (Palop and Mucke, 2016; Schmid *et al*., 2016; Ambrad Giovannetti and Fuhrmann, 2019; Xu *et al*., 2020). All extrinsic and intrinsic simulation features of the three scenarios of hyperexcitability discussed here can be seen in **Supplementary Table S4**.

### Ion channel modifications

AD-related changes of ion channels were modeled by modifying their respective conductances according to experimental reports. To reduce ion channel expression or increase its density, the maximum channel conductance was scaled by a corresponding factor. Specifically, the voltage-dependent sodium channel *INa* in the axon (Liu *et al*., 2015; Wang *et al*., 2016; Ghatak *et al*., 2019) was upscaled 1.3-fold and the persistent sodium channel *INap* in the soma (Williams and Stuart, 1999; Yue *et al*., 2005; Beck and Yaari, 2008) was upscaled 2-fold in **Figure 4E** of Scenario 3 for extrinsic (*E/I* balance) and intrinsic (ion channel) modifications. At the same time the medium afterhyperpolarisation calcium-activated potassium channel *I_AHP_* in the soma and apical dendrites was downscaled 0.85-fold (Beck and Yaari, 2008; Zhang *et al*., 2014; Wang *et al*., 2015b,a; Niday and Bean, 2021) in **Figure 4E**, while the T-type calcium channel in dendrites (Yaari *et al*., 2007; Beck and Yaari, 2008; Cain and Snutch, 2013; Medlock *et al*., 2018; Garg *et al*., 2021) was upscaled 2-fold. For the effect of ion channel changes alone in **Supplementary Figure S4C** we scaled the aforementioned ion channel conductances to additional values (0.05 for *I_AHP_* and 3 for *I_Nap_*, *I_Na_* and *I_CaT_*) in order to investigate a possible maximum burst firing increase (with minimal change in single spike firing) under pathological ion channel conditions. Furthermore, we tested alterations of other ion channels that are mentioned in literature in relation to AD including potassium and hyperpolarisation-activated, cyclic nucleotide-gated *HCN* channels, which have been proposed to be involved in hyperexcitability (Beck and Yaari, 2008; Musial *et al*., 2018; Vitale *et al*., 2021). However, our simulations did not show a significant contribution to the increase in burst activity of APP/PS1 CA1 PC morphologies. Modifications in the conductance of the L-type calcium channel did not enhance burst firing as well (Anekonda *et al*., 2011; Berridge, 2014).

### Statistical Analysis

We used an unpaired Student’s t-test for the statistical comparison of morphological parameters (path length, number of branchpoints, surface area, dendritic diameter), voltage change and firing rate in WT and APP/PS1 groups in **Figures 1 - 3** and **Supplementary Figures S2 and S3**. When we assessed the statistical significance of firing rate, event rate (singlets, bursts, triplets) and initial firing rate between the WT and APP/PS1 groups as well as the APP/PS1 groups with modified extrinsic and intrinsic properties in **Figure 4** and **Supplementary Figures S4, S5 and S6** (with additional WT groups of modified extrinsic and intrinsic properties) we used the one-way ANOVA. All error bars are shown with mean and standard error of mean. A p-value of *p <* 0.05 was considered significant and depicted with an asterisk.

## Additional information

### Competing interests

The authors declare to have no competing financial interests.

## Funding

This work was supported by BMBF (No. 01GQ1406 - Bernstein Award 2013 to HC, No. 031L0229, to PJ), by University Medical Center Giessen and Marburg (UKGM; to PJ) and by funds from the von Behring Rontgen Foundation (to PJ).

---

**Figure S1.**
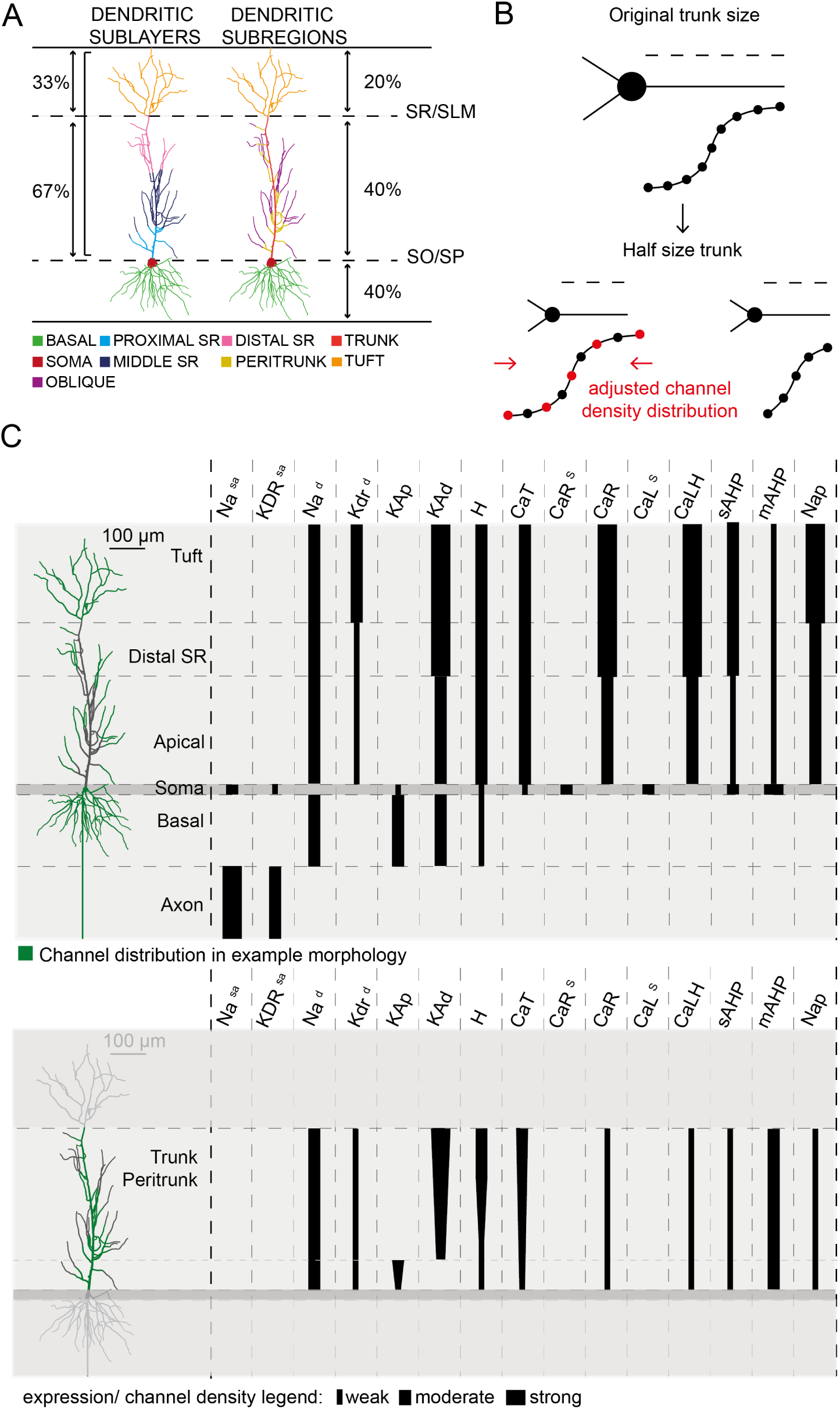
Dendritic regions and ion channel distributions of the model by Poirazi *et al*. (2003b) in reconstructed CA1 pyramidal cells. **A**, Dendritic layer distribution: Each subdivision represents the estimated boundary for changes in electrotonic properties and channel densities based on the dendritic length contribution that reflects the absolute distance inflexion points employed in the compartmental model. Dendritic regions distribution includes the “peritrunk area” according to Poirazi *et al*. (2003b), which constitutes the first 50*µm* of each oblique branch stemming away from the apical trunk. **B**, Schematic of the rescaling process for the nonlinear distribution of ion channels in morphologically diverse CA1 pyramidal cell models. *Top*: original cell morphology’s trunk laminar depth, from 0 to 8 *µ*m, in steps of 1. *Bottom left*: cell with an apical trunk half the size of the original. The original laminar depth vector, is resampled, taking steps double the size of the original ones (size 2). *Bottom right*: the vector is then compressed to fit the new cell’s trunk size. **C**, Schematic summary of the ion channel composition of the CA1 pyramidal cell model with Poirazi biophysics (Poirazi *et al*., 2003b). *Left*: an exemplary morphology out of all reconstructed cell morphologies used for the compartmental modelling. *Top right*: Distribution of ion channel density for all regions besides the apical trunk and peritrunk areas. *Bottom right*: Distribution of ion channel density for the apical trunk and peritrunk areas. The relative thickness of the lines indicates the channel density (non-existent, weak, moderate or strong), and the shape depicts the uniformity of their spatial distribution (uniform or linearly increasing).

**Figure S2.**
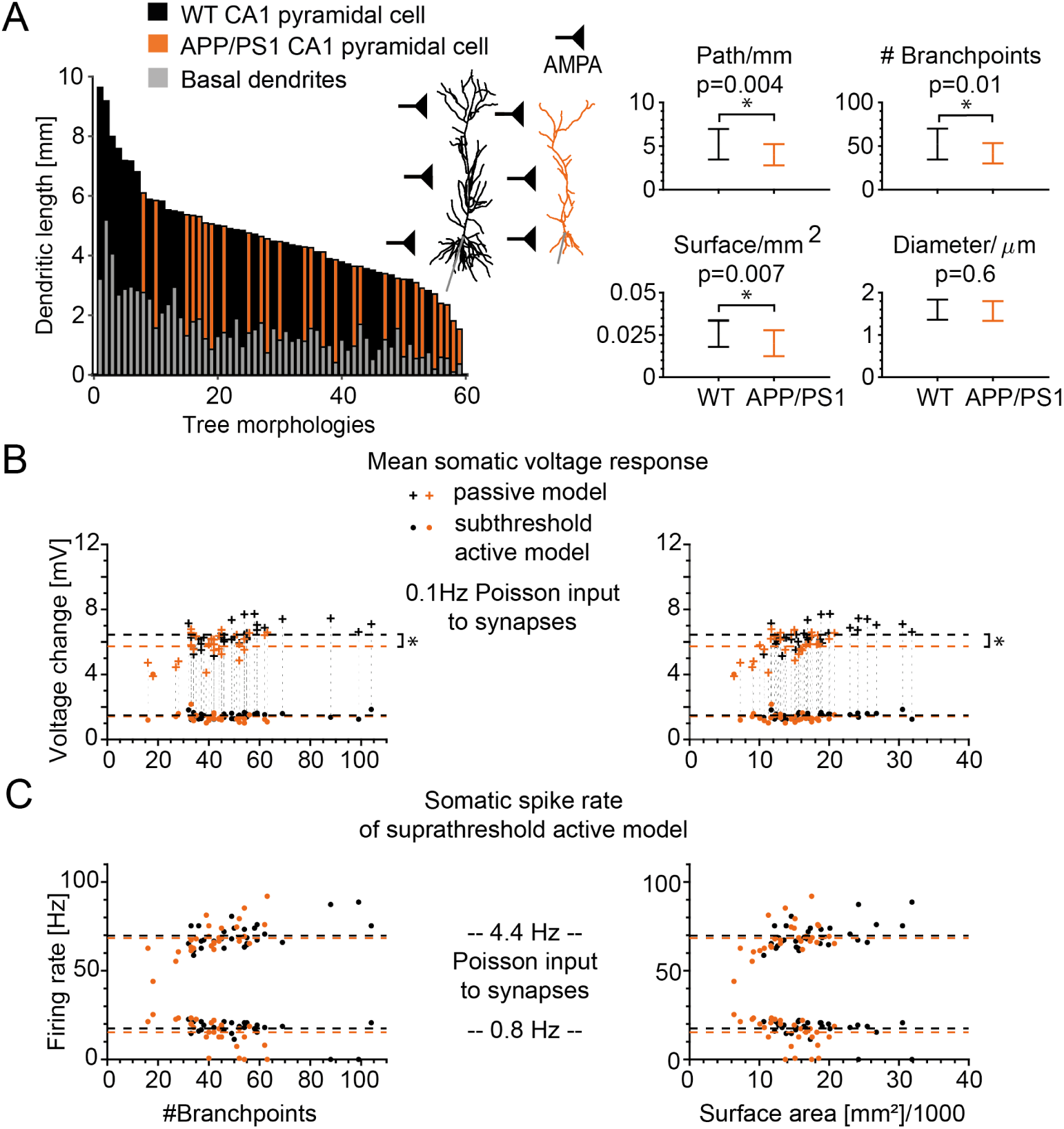
Overview of morphological measures for reconstructed WT and APP/PS1 CA1 cells and their influence on responses to distributed synaptic inputs. **A**, *Left*: Sorted distribution of the dendritic length of all cells shown, ranked from longest to shortest. *Black* denotes the WT cell, *orange* the APP/PS1 cell and *grey* is the basal portion of the total dendritic length. *Right*: The comparison of average dendritic path length (WT 5.19 *±* 1.75*mm*, APP/PS1 4.00 *±* 1.22*mm*, *p* = 0.004), number of branching points (WT 52.16 *±* 17.79, APP/PS1 41.61 *±* 11.67, *p* = 0.01) and dendrite surface area (WT 0.026 *±* 0.008*mm*^2^, APP/PS1 0.02 *±* 0.008*mm*^2^, *p* = 0.007) between WT and APP/PS1 model cell groups shows significant differences. The average dendritic diameter is similar in both cell groups (WT 1.6 *±* 0.24*µm*, APP/PS1 1.57 *±* 0.23*µm*, *p* = 0.6). As part of the morphologies preprocessing the diameter was subsequently normalised to *d* = 1*µm*. **B**, Voltage change responses of passive and subthreshold active cells to distributed AMPA inputs as in Figure 1B but plotted against the number of branchpoints (*left*) and the surface area (*right*). The dashed lines show the mean activity of the WT (*black*) and APP/PS1 (*orange*) CA1 cell groups (mean voltage passive model: WT 6.46 *±* 0.67*mV*, APP/PS1 5.74 *±* 0.80*mV*, the asterisk depicts *p <* 0.0005; mean voltage subthreshold active model: WT 1.47 *±* 0.17*mV*, APP/PS1 1.43 *±* 0.56*mV*). **C**, Firing rate responses to active distributed AMPA inputs as in Figure 1D but plotted against the number of branchpoints (*left*) and the surface area (*right*). The dashed lines show the mean firing rate (0.8*Hz* input: WT 17.42 *±* 5.28*Hz*, APP/PS1 15.43 *±* 7.75*Hz*; 4.4*Hz* input: WT 69.89 *±* 7.00*Hz*, APP/PS1 68.40 *±* 9.37*Hz*).

**Figure S3.**
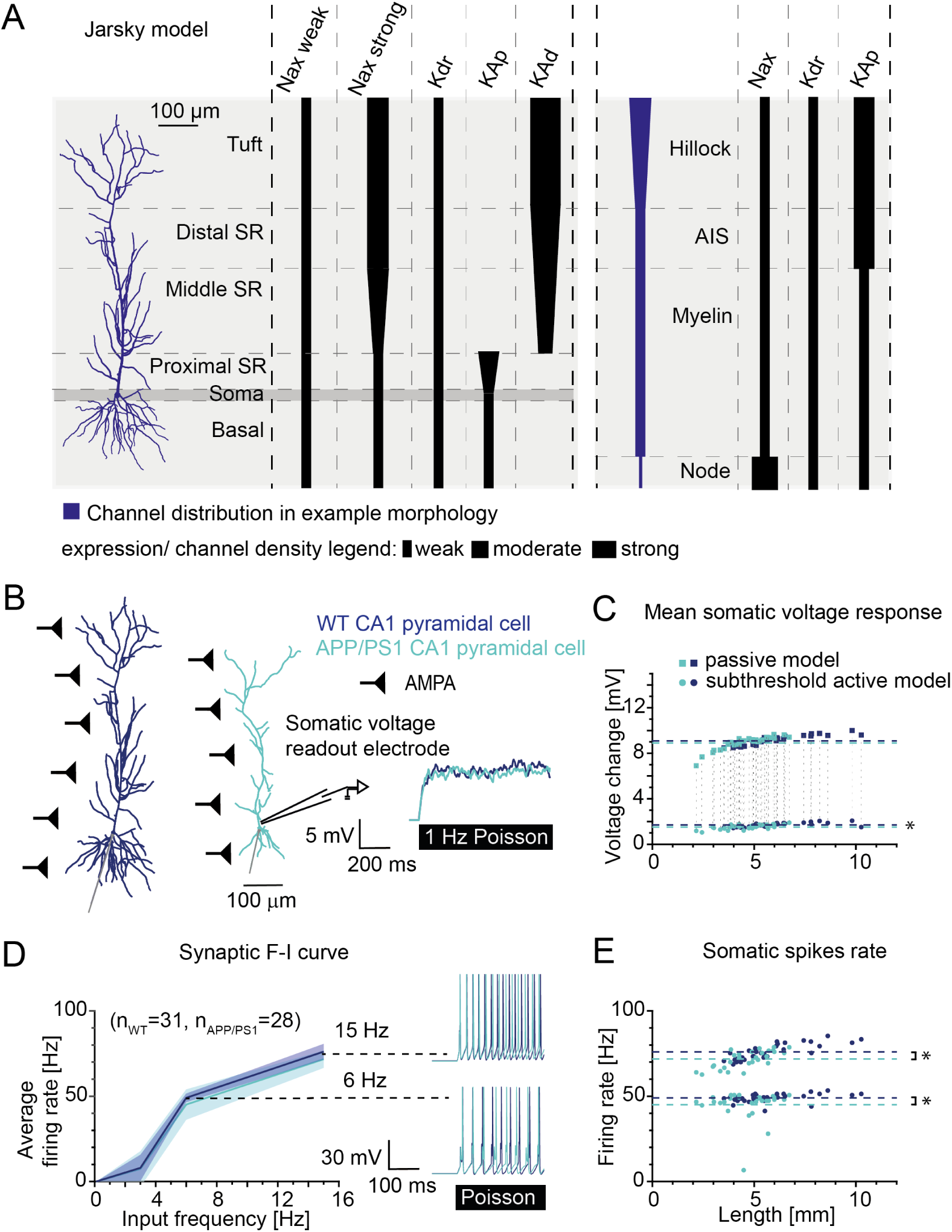
Responses to distributed synaptic inputs show no hyperexcitability in APP/PS1 morphologies with the biophysical model by Jarsky *et al*. (2005) **A**, Schematic summary of the ion channel composition of the CA1 pyramidal cell model with biophysics by Jarsky *et al*. (2005). *Left*: An exemplary morphology out of all reconstructed cell morphologies used for the compartmental modelling. *Right*: Distribution of ion channel density for all regions. The relative thickness of the lines indicates the channel density (non-existent, weak, moderate or strong), and the shape depicts the uniformity of their spatial distribution (uniform or linearly increasing). **B**, *Left*: Sample 3D-reconstructed morphologies of CA1 pyramidal cells (WT in *dark blue*, APP/PS1 in *light blue*) with a schematic of homogeneously distributed AMPA synapses with Poisson input pattern such as in Figure 1A. *Right*: Sample trajectories for the voltage response of the two sample cells for the passive model with whole cell distributed AMPA stimulation of 1*Hz* Poisson inputs. **C**, Voltage change responses to distributed AMPA inputs at 1*Hz* for the passive (*squares*) and 0.5*Hz* for the subthreshold active model by Jarsky *et al*. (2005, *circles*) for all available cell morphologies (WT: *n* = 31, APP/PS1: *n* = 28). The dashed lines show the mean activity of the WT (*dark blue*) and APP/PS1 (*light blue*) CA1 cell groups (mean voltage passive model: WT 9.11 *±* 0.43*mV*, APP/PS1 8.91 *±* 0.64*mV*; mean voltage subthreshold active model: WT 1.69 *±* 0.18*mV*, APP/PS1 1.53 *±* 0.22*mV*, the asterisk depicts *p* = 0.0036). **D**, Synaptic input-output firing curve for the somatic firing rate of WT and APP/PS1 pyramidal cells in active compartmental models with AMPA synapses. The input frequency ranges from 0.5*Hz* to 15*Hz*. The *right* insets show AP firing traces of the two sample cells for an input frequency of 6*Hz* and 15*Hz* respectively. **E**, Firing rate versus dendritic length corresponding to the data points with input frequency of 6*Hz* and 15*Hz* in **D**. The dashed lines show the mean firing rates (6*Hz* input: WT 49.05 *±* 2.52*Hz*, APP/PS1 45.12 *±* 8.9*Hz*, *p* = 0.0217; 15*Hz* input: WT 76.11 *±* 4.58*Hz*, APP/PS1 71.66 *±* 5.07*Hz*, *p <* 0.0009). For all simulations the AMPA synapse strength is 0.1*nS* and the density is homogeneous with 1 synapse per 2*µm*.

**Figure S4.**
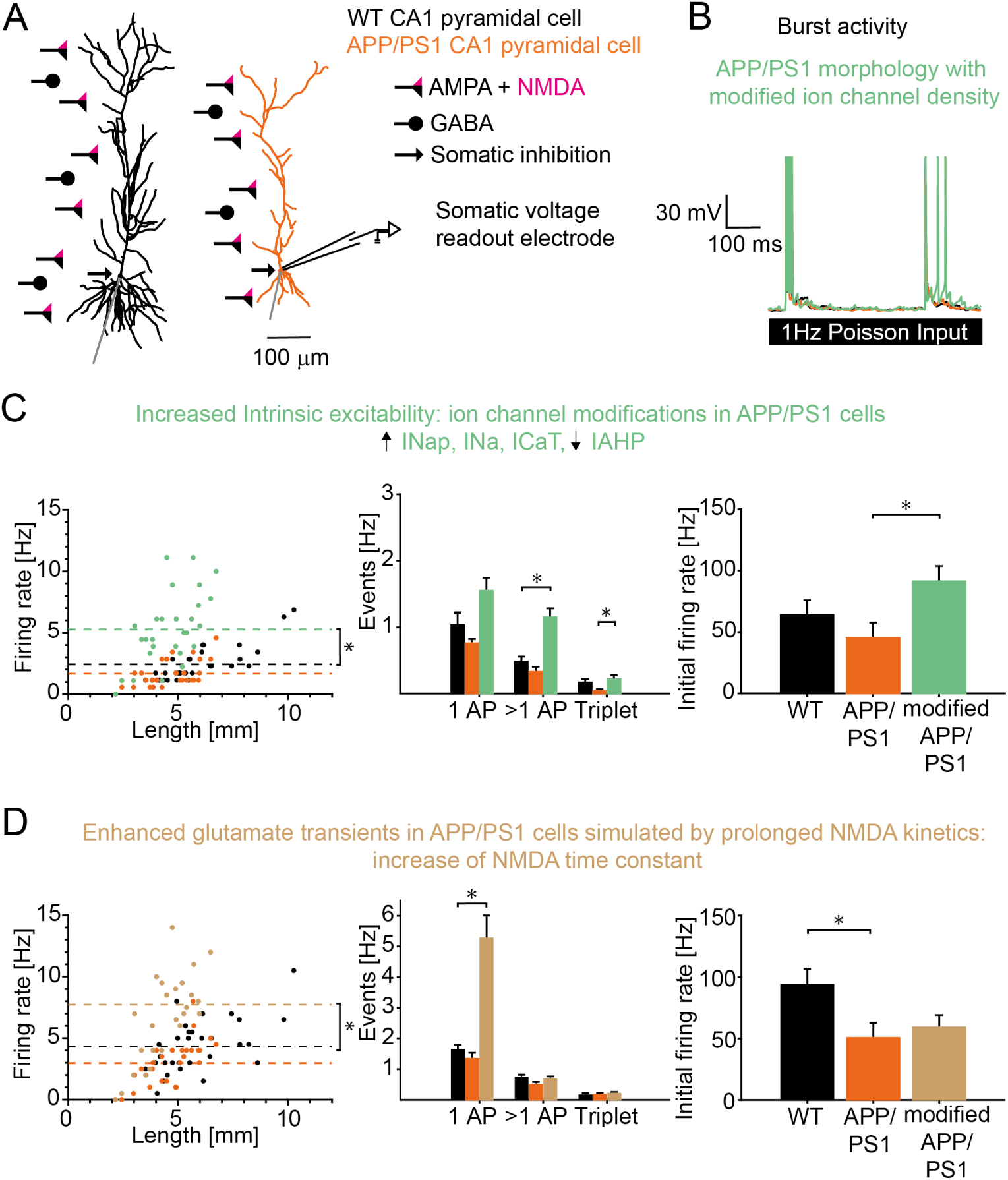
Qualitative but not quantitative reproduction of increased burst firing in APP/PS1 cells due to intrinsic ion channel changes and failed reproduction of increased burst firing due to enhanced glutamate transients simulated as prolonged NMDA kinetics. **A**, Sample morphologies of CA1 pyramidal cells (WT in *black*, APP/PS1 in *orange*) with a schematic of distributed AMPA (*black* bar and triangle), NMDA (*magenta* triangle) and GABA (*black* bar and circle) synapses (**Supplementary Table S3**) such as in Figure 4A. **B**, Sample trajectories for the voltage response of the two sample cells in **A**. A third group was added with the same APP/PS1 morphologies but additionally modified intrinsic ion channels (*light green*). The traces correspond to the whole cell distributed synapse stimulation of 1*Hz* Poisson input. **C**, *Left*, Firing rate versus dendritic length of WT, APP/PS1 and APP/PS1 cells with intrinsic ion channel changes (*light green*: increased *I_Nap_*, *I_Na_*, *I_CaT_* densities, decreased *I_AHP_* density; see details in **Methods**). The dashed lines indicate the mean firing rate. The asterisk depicts a significant firing rate increase with *p <* 1 *·* 10*^−^*^6^. *Middle*, Number of events for the same cell groups: single AP, bursts (*ISI ≤* 13.3*ms*) and triplets. The asterisk indicates *p <* 2 *·* 10*^−^*^5^ for an increased burst rate between WT and modified APP/PS1 with ion channel alterations. The number of triplets is increased as well between APP/PS1 and modified APP/PS1 with ion channel alterations (asterisk depicts *p* = 0.04), but not significantly between WT and modified APP/PS1 with ion channel alterations (*p* = 0.73). Note that experiments (Šišková *et al*., 2014) showed unchanged 1*AP* but strongly increased bursts and triplet AP firing in APP/PS1 pyramidal neurons. *Right*, Initial firing rate of the first two APs (*p* = 0.03). **D**, *Left*, Firing rate versus dendritic length of WT, APP/PS1 and APP/PS1 cells with the enhancement of extrinsic glutamate transients simulated as increased NMDA decay time constant: *τ_enhancedNMDA_*= 2 *· τ_W T,NMDA_*. The dashed lines indicate the mean firing rate. The asterisk depicts *p* = 0.0002. *Middle*, Number of events for the same cell groups: single AP, bursts and triplets. The asterisk indicates *p <* 3 *·* 10*^−^*^7^ for an increased single spike rate between WT and modified APP/PS1 with increased NMDA decay. *Right*, Initial firing rate of the first two APs (*p <* 0.03).

**Figure S5.**
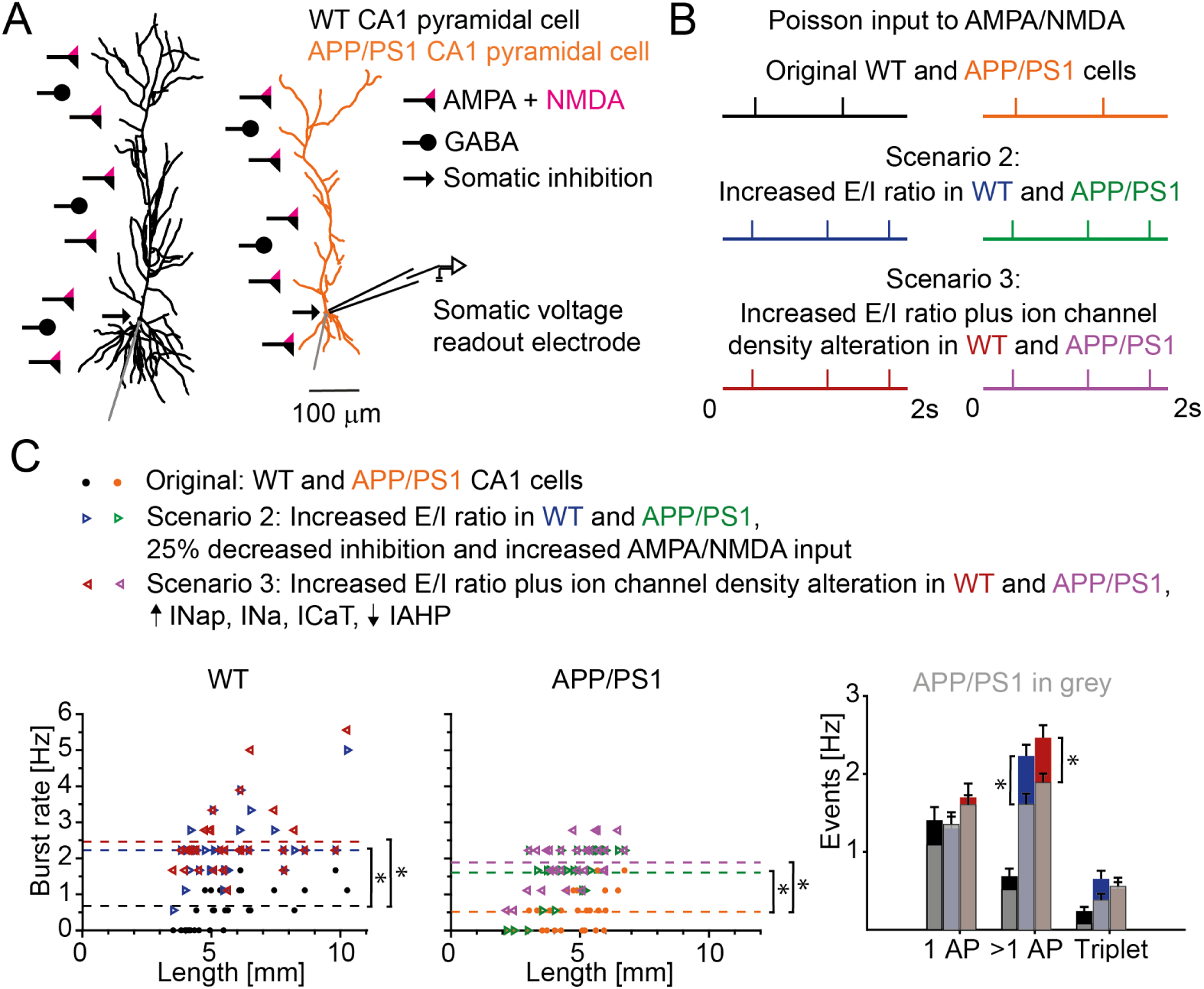
Control simulations in WT morphologies modified by identical extrinsic (network) and intrinsic (ion channel) changes as in modified APP/PS1 morphologies lead to a stronger transition from solitary to burst firing in WT than in APP/PS1 morphologies. **A**, Sample morphologies of CA1 pyramidal cells (WT in *black*, APP/PS1 in *orange*) with a schematic of heterogeneously distributed AMPA (*black* bar and triangle), NMDA (*magenta* triangle) and GABA (*black* bar and circle) synapses (**Supplementary Table S3**) such as in Figure 4A. **B**, Example Poisson input pattern with 1*Hz* frequency to AMPA/NMDA synapses of original WT (*black*) and APP/PS1 cells (*orange*), and input pattern with 1.3*Hz* frequency for WT (*blue*) and APP/PS1 cells (*green*) with increased *E/I* ratio, and for WT (*red*) and APP/PS1 (*purple*) with increased *E/I* ratio plus ion channel alterations. **C**, *Left*: Burst firing rate of the original WT morphologies with no additional extrinsic/intrinsic changes (*black circle*) and increased burst firing of the modified WT morphologies (*blue* and *red triangles*) depending on their dendritic length. The dashed lines indicate the mean firing rate. The asterisks depict *p* = 2.1 *·* 10*^−^*^8^ for original WT and both modified WT groups. *Middle*: Burst firing rate of the original APP/PS1 morphologies (*orange circle*) and increased burst firing of the modified APP/PS1 morphologies (*green* and *purple triangles*) depending on their dendritic length. The dashed lines indicate the mean firing rate. The asterisks depict *p* = 4 *·* 10*^−^*^7^ for original APP/PS1 and modified APP/PS1 with extrinsic changes, and *p* = 2.1 *·* 10*^−^*^8^ for original APP/PS1 and modified APP/PS1 with extrinsic/intrinsic changes. *Right*: Modified WT cells (*blue* and *red*) show a stronger increase in burst firing (extrinsic changes: *p* = 0.0164, extrinsic/intrinsic changes: *p* = 0.034) than modified APP/PS1 cells (*grey*).

**Figure S6.**
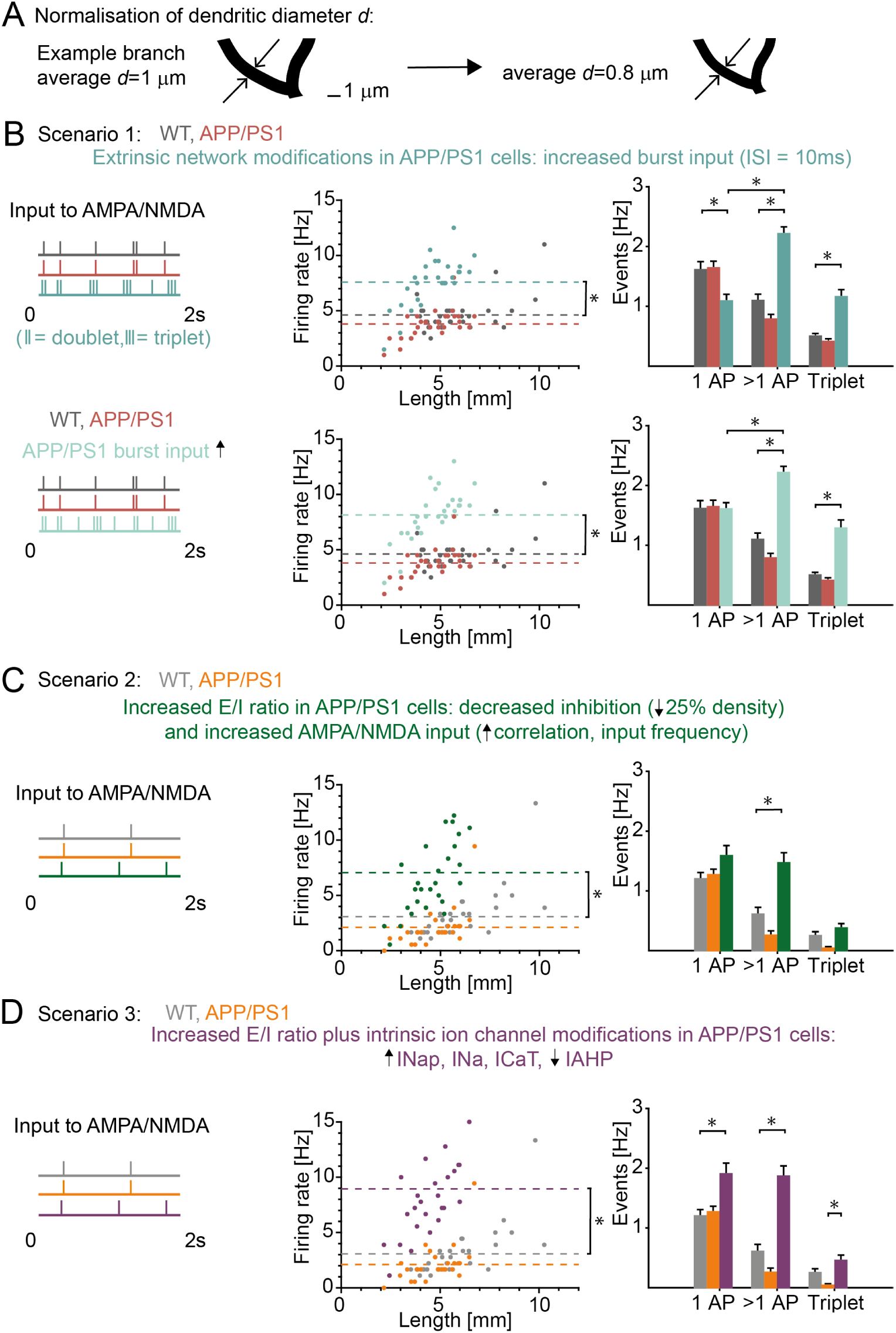
Control simulations of morphologies with normalised average dendritic diameter of *d* = 0.8*µm* modified by identical extrinsic (network) and intrinsic (ion channel) changes as in morphologies with *d* = 1*µm* lead to an increase in burst firing in APP/PS1 morphologies. **A**, *Left*, Example branch of a reconstructed CA1 pyramidal cell with normalised average dendritic diameter. The arrows indicate the adopted diameter for a specific compartment of the reconstructed cell to guarantee an overall average target diameter of the whole dendritic morphology of *d* = 1*µm*. *Right*, The normalised average dendritic diameter was reduced to *d* = 0.8*µm* according to Benavides-Piccione *et al*. (2020). **B**, Scenario 1: *Top*, Similar to Figure 4 **B, C** but with normalised dendritic diameter of *d* = 0.8*µm*. The asterisk for the overall firing rate depicts *p <* 3 *·* 10*^−^*^8^. The asterisks for the specific event rate indicate *p* = 0.008 for singlets, *p* = 1 *·* 10*^−^*^9^ for bursts and *p <* 7 *·* 10*^−^*^8^ for triplets respectively. The mode change from single spikes to predominantly bursts is significant with *p <* 5 *·* 10*^−^*^9^. Note that experiments (Šišková *et al*., 2014) showed unchanged single spikes but strongly increased number of bursts (*>* 1*AP*) and increased triplet AP firing in APP/PS1 pyramidal neurons. Here the burst increase was even stronger and the single AP rate decreased. *Bottom*, Similar to **Top** but with changed input pattern (*Left*) to get firing and burst rate as in Figure 4 **C**. The asterisk for the overall firing rate depicts *p <* 2 *·* 10*^−^*^9^. The asterisks for the specific event rate indicate *p <* 1 *·* 10*^−^*^9^ for bursts and *p <* 2 *·* 10*^−^*^8^ for triplets respectively. The mode change from single spikes to predominantly bursts is significant with *p <* 0.0002. **C**, Scenario 2: Similar to Figure 4 **B, D** but with normalised dendritic diameter of *d* = 0.8*µm*. The asterisk for the overall firing rate depicts *p* = 5 *·* 10*^−^*^6^. The asterisk for the specific event rate indicates *p <* 2 *·* 10*^−^*^5^ for bursts. **D**, Scenario 3: Similar to Figure 4 **B, E** but with normalised dendritic diameter of *d* = 0.8*µm*. The asterisk for the overall firing rate depicts *p <* 2 *·* 10*^−^*^9^. The asterisks for the specific event rate indicate *p <* 0.0008 for singlets, *p <* 3 *·* 10*^−^*^9^ for bursts and *p <* 0.0002 for triplets respectively.

**Table S1.** Spiking features of the model by Poirazi *et al*. (2003b)

| Spiking features | Original | T2N version | WT | APP/PS1 |
| --- | --- | --- | --- | --- |
| Input resistance | 65.25MΩ | 64.16MΩ | 117.04 ± 22.11MΩ | 133.70 ± 33.15MΩ |
| Total Capacitance | 115.60pF | 110.88pF | 96.92 ± 31.55pF | 75.85 ± 20.42pF |
| Membrane Time Constant | 11.02ms | 12.68ms | 9.29 ± 1.10ms | 8.34 ± 0.90ms |
| Resting Potential | -68.40mV | -68.40mV | -68.39 ± 0.20mV | -68.40 ± 0.34mV |
| Spike Threshold | -58.60mV | -58.51mV | -60.07 ± 0.34mV | -59.99 ± 0.16mV |
| Spike Amplitude | 80.00mV | 77.88mV | 81.04 ± 3.02mV | 82.25 ± 3.26mV |
| Spike Peak | 21.11mV | 19.33mV | 20.97 ± 3.05mV | 22.25 ± 3.30mV |
| Spike Width (at -30mV) | 1.77ms | 1.75ms | 1.46 ± 0.11ms | 1.43 ± 0.06ms |
| AHP<sup>1</sup> Amplitude | 1.67mV | 1.30mV | 1.40 ± 0.20mV | 1.31 ± 0.26mV |
| Rheobase | 120pA | 130pA | 33 ± 14pA | 28 ± 12pA |

**Table S2.** Spiking features of the model by Jarsky *et al*. (2005)

| Spiking features | Original | T2N version | WT | APP/PS1 |
| --- | --- | --- | --- | --- |
| Input resistance | 67.72MΩ | 68.46MΩ | 212.05 ± 60.74MΩ | 262.68 ± 94.45MΩ |
| Total Capacitance | 81.84pF | 80.93pF | 54.22 ± 18.88pF | 41.68 ± 12.37pF |
| Membrane Time Constant | 7.74ms | 7.59ms | 12.51 ± 1.30ms | 12.90 ± 1.92ms |
| Resting Potential | -71.95mV | -72.25mV | -71.38 ± 0.29mV | -71.38 ± 0.47mV |
| Spike Threshold | -62.92mV | -62.30mV | -53.80 ± 5.30mV | -55.57 ± 5.98mV |
| Spike Amplitude | 97.93mV | 97.51mV | 89.56 ± 5.75mV | 91.37 ± 6.11mV |
| Spike Peak | 34.93mV | 35.05mV | 35.75 ± 1.66mV | 35.80 ± 1.17mV |
| Spike Width (at -30mV) | 0.82ms | 0.82ms | 0.79 ± 0.05ms | 0.79 ± 0.04ms |
| AHP<sup>1</sup> Amplitude | 5.80mV | 7.28mV | 17.05 ± 5.83mV | 14.91 ± 6.90mV |
| Rheobase | 130pA | 150pA | 119 ± 38pA | 95 ± 35pA |

**Table S3.** Modeled synapse densities based on data from Megías *et al*. (2001), Bloss *et al*. (2016) and Šišková *et al*. (2014)

| Dendritic region | AMPA,NMDA synapse/μm | GABA synapse/μm |
| --- | --- | --- |
| Proximal basal | 0.1 | 0.3 |
| Middle basal | 0.3 | 0.2 |
| Distal basal | 2.0 | 0.06 |
| Proximal apical trunk | 0.3 | 0.3 |
| Distal apical trunk | 3.5 | 0.06 |
| Proximal apical oblique | 2.2 | 0.05 |
| Distal apical oblique | 2.7 | 0.05 |
| Proximal tuft | 1.2 | 0.2 |
| Distal tuft | 0.8 | 0.2 |

For APP/PS1 morphologies the AMPA synapse density is equal to WT morphologies in all dendritic regions except for the tuft region where it reduces to 82.3% of WT synapse density. This is based on spine density data (Šišková *et al.*, 2014).

**Table S4.** Extrinsic and intrinsic simulation details for the three scenarios of hyperexcitability in AD

| Simulation features | Scenario 1 control | Scenario 1 increased burst input | Scenario 2/3 control | Scenario 2 increased $E/I$ | Scenario 3 channel alterations |
| --- | --- | --- | --- | --- | --- |
| input frequency | $3Hz$ | $7Hz$ | $1Hz$ Poisson | $1.3Hz$ Poisson | $1.3Hz$ Poisson |
| input burst frequency | $0.5Hz$ | $2.5Hz$ | 0 | 0 | 0 |
| input cv | 0 | 0 | 0.41 | 0.17 | 0.17 |
| input correlation | 1 | 1 | 0.4 | 0.8 | 0.8 |
| % AMPA activation | 4 | 4 | 100 | 100 | 100 |
| % GABA activation | 100 | 100 | 100 | 75 | 75 |
| scale $I_{Na}$ conductance | 1 | 1 | 1 | 1 | 1.3 |
| scale $I_{Nap}$ conductance | 1 | 1 | 1 | 1 | 2 |
| scale $I_{CaT}$ conductance | 1 | 1 | 1 | 1 | 2 |
| scale $I_{AHP}$ conductance | 1 | 1 | 1 | 1 | 0.85 |

## Notes

### Competing Interest Statement

The authors have declared no competing interest.

